# Discovery of free glycated amines and glycated urea as potential markers of diabetic nephropathy

**DOI:** 10.1101/2024.02.17.580794

**Authors:** Rashdajabeen Q Shaikh, Sancharini Das, Arvindkumar Chaurasiya, MG Ashtamy, Amreen B Sheikh, Moneesha Fernandes, Shalbha Tiwari, AG Unnikrishnan, Mahesh J Kulkarni

**Affiliations:** Biochemical Sciences Division, CSIR-National Chemical Laboratory, Pune- 411008, India; Academy of Scientific and Innovative Research (AcSIR), Ghaziabad, UP-201002, India; Organic Chemistry Division, CSIR-National Chemical Laboratory, Pune-411008, India; Department of Diabetes and Endocrine Research, Chellaram Diabetes Institute, Pune-411021, India

**Keywords:** Glycation, mass spectrometry, metabolomics, amino acids, biomarker, kidney

## Abstract

The role of protein glycation in the pathogenesis of diabetes is well established. Akin to proteins, free amino acids and other small-molecule amines are also susceptible to glycation in hyperglycemic conditions and may have a role in the pathogenesis of the disease. However, the information on glycation of free amino acids and other small-molecule amines is relatively obscure. In the quest to discover small molecule glycated amines in the plasma, we have synthesized glycated amino acids, glycated creatine, and glycated urea, and by using a high-resolution accurate mass spectrometer, a mass spectral library was developed comprising the precursor and predominant fragment masses of glycated amines. Using this information, we report the discovery of glycation of free lysine, arginine, and leucine/isoleucine from the plasma of diabetic patients. This has great physiological significance as glycation of these amino acids may create their deficiency and affect vital physiological processes such as protein synthesis, cell signaling and insulin secretion. Also, these glycated amino acids could serve as potential markers of diabetes and its complications. On the other hand, accumulation of creatinine and urea in the plasma act as biomarkers of diabetic nephropathy diagnosis. For the first time, we report the detection of glycated urea in diabetes, which is confirmed by matching the precursor and fragment masses with the *in vitro* synthesized glycated urea by using ^12^C6 and ^13^C6-glucose. Further, we quantified glycated urea detected in two forms, monoglycated urea (MGU) and diglycated urea (DGU), by a targeted mass spectrometric approach in the plasma of healthy, diabetes, and diabetic nephropathy subjects. Both MGU and DGU showed a positive correlation with clinical parameters such as blood glucose and HbA1c. Given that urea gets converted to glycated urea in hyperglycemic conditions, it is crucial to quantify MGU and DGU along with the urea for the diagnosis of diabetic nephropathy.

## Introduction

Protein glycation, a non-enzymatic reaction between reducing sugars and proteins, has been a well-studied process in diabetes [1]. This reaction was first reported by Louis Camille Maillard in 1912 while heating amino acids and reducing sugars. Thus, it is also referred to as Maillard’s reaction. Since then, glycation has been studied; however, it is mainly limited to proteins [2]. Especially, the glycation of abundant large molecular weight proteins such as hemoglobin and albumin is documented in great detail by various researchers [3]. Glycated hemoglobin (HbA1c) is used as a diagnostic marker for diabetes management [4]. Glycated albumin, on the other hand, acts as a predominant advanced glycation end product (AGE) in circulation and is known to have harmful physiological consequences through interaction with the receptor for AGE (RAGE) [5]. Both these proteins are abundant and long-lived; thus, they are more prone to undergo glycation. In contrast, the low-abundant proteins with a shorter lifespan are less likely to get glycated, although a few low-abundant proteins, such as insulin, hexokinase, catalase, superoxide dismutase (SOD), and glutathione peroxidase, are found to be glycated [6]. The impact of glycation on such low-abundant protein can not be undermined. For example, *in vitro*, glycated insulin has a lesser binding affinity for insulin receptors and interacts with RAGE to activate oxidative stress and pro-inflammatory pathways [7]. Similarly, free amino acids are also likely to undergo glycation. In yeast, the formation of glycated products such as pyrraline, formyline, and maltosine from free amino acids was observed during the brewing of beer [8]. In diabetes, there is a possibility of free amino acids and other amines undergoing glycation in hyperglycemic conditions. However, there are no studies in the literature with respect to the glycation of free amino acids and small molecule amines. This could be due to a lower concentration, shorter half-life, and a lack of methods or libraries to identify glycated amines. In light of this, we have made an attempt to discover novel small-molecule glycated amines from diabetic plasma. To this end, we have synthesized glycated forms of amino acids, urea, and creatine, and using the information on precursor and fragment masses, discovered novel glycated amines and quantified glycated urea in the plasma of healthy (H), diabetes (DM) and diabetic nephropathy (DN) subjects by using a targeted metabolomics approach.

## 2. Materials and methods

### Materials

Reagents, such as LC-MS-grade water, methanol (MeOH), and acetonitrile (ACN), were purchased from JT Baker (PA, USA). Formic acid, Sodium Phosphate Dibasic (Na2HPO4), Sodium Phosphate Monobasic (NaH2PO4), all 20 amino acid standards *viz*., alanine, arginine, asparagine, aspartic acid, cysteine, glutamic acid, glutamine, glycine, histidine, isoleucine, leucine, lysine, methionine, phenylalanine, proline, serine, threonine, tryptophan, tyrosine and valine, urea, creatine and glucose were obtained from Sigma-Aldrich (St. Louis, MO, USA). Thermo Hypersil Gold C18 column (length 150 mm, ID 2.1 mm, particle size 1.9 µM and pore size 175 Å) was procured from ThermoFisher Scientific (Lithuania, Europe).

### 2.1. Study design

This study was carried out to discover novel small molecule glycated amines in the plasma. Glycated amino acids, glycated creatine, and glycated urea were synthesized *in vitro* to identify glycated amines. The mass spectral library comprising precursor and product masses was developed for *in vitro* synthesized glycated amines. Using this information, glycated amines from the plasma of diabetic patients were identified. The plasma samples used in this study were collected in a previous study to discover risk prediction markers for diabetic nephropathy from Chellaram Diabetes Institute, Pune, India, with the approval of the Institutional Ethics Committee. A signed written informed consent was obtained from all subjects before the blood collection. The procedure for plasma collection was detailed earlier [9].

### 2.2 *In vitro* synthesis of glycated amino acids, urea, and creatine

The glycation reaction was carried out by incubating 500 µl of each amino acid (100 mM) *viz*., alanine, arginine, asparagine, aspartic acid, cysteine, glutamic acid, glutamine, glycine, histidine, isoleucine, lysine, methionine, proline, serine, threonine, and valine with 500 µl of D-glucose (500 mM) in 50 mM phosphate buffer (pH 7.4) at 37oC for 72 h. The concentration was 50 mM for leucine, while for tyrosine, tryptophan, and phenylalanine, it was 3.125 mM, and proportionately, glucose concentration was diluted. Similarly, glycated creatine and glycated urea were synthesized by incubating 500 µl of creatine (25 mM) and urea (100 mM) with 500 µl of glucose (500 mM). Further, the *in vitro* synthesized glycated amino acids, glycated creatine, and glycated urea were analyzed by parallel reaction monitoring using LC-HRMS (Thermo Q-Exactive Orbitrap mass spectrometer) studies followed by XIC-based confirmation.

### 2.3. Extraction of metabolites from plasma samples

For the extraction of metabolites, 100 µl of plasma was mixed with 400 µl of chilled methanol and incubated for 1 h at -20℃. The precipitated protein was separated by centrifugation for 20 min at 13000 g, and the supernatant containing metabolites was dried by using a vacuum concentrator at 4oC. The sample was reconstituted in 100 µl of 50% methanol, sonicated for 2 min, centrifuged for 15 min, and used for mass spectrometric analysis [10]. The extracted metabolites were analyzed to detect glycated amines by parallel reaction monitoring as described above for *in vitro* glycated amines.

### 2.4. LC-HRMS analyses

The synthesized glycated amines were acquired in parallel reaction monitoring (PRM) using a UHPLC-High-Resolution Accurate mass spectrometer (LC-HRAM, Orbitrap, ThermoFisher Scientific) equipped with a heated electrospray ionization (HESI) source. The mass spectrometer was calibrated before acquisition. Individual glycated amine (1µl) was loaded onto the Thermo Hypersil Gold C18 column (length 150 mm, ID 2.1 mm, particle size 1.9 µM, and pore size 175 Å). The column was maintained at 40°C throughout the acquisition time. Glycated amines are eluted from the column in an isocratic mode through the mobile phase consisting of 1:1 acetonitrile and water, supplemented with 0.1% formic acid at a flow rate of 200 µl/min with 5 min runtime [11].

The precursor masses to be detected were specified in the inclusion list of the PRM method. The precursor ions were fragmented at a normalized collision energy of 30eV, and the fragments were acquired at a resolution of 17500. AGC target was set to 2e5, maximum IT to 100 ms, and isolation window to 4.0 m/z. Sheath gas and auxiliary gas flow rates were set to 40 and 15, respectively.

### 2.5. Screening and identification of glycated amines from the plasma

A list of accurate masses of glycated amino acids, glycated creatine, and glycated urea is tabulated (Table 1). The presence of these masses was investigated by an extracted ion chromatogram. The glycated amines are confirmed by matching the precursor and fragment ions with the *in vitro* synthesized glycated amines (Table 2).

**Table 1:** HRMS analysis of *in vitro* synthesized glycated amines with most intense fragment ions.

| S. No. | Glycosylated amines | Monoisotopic mass (m/z) | Observed mass (m/z) | RT (min) | Fragment ions | Intensity |
| --- | --- | --- | --- | --- | --- | --- |
| 1 | Monoglycosylated arginine | 337.1723 | 337.1707 | 1.53 | 70.0654<br>112.0869<br>114.1026<br>173.1393<br>175.1185<br>217.1287 | 1.53e6 |
| 2 | Diglycosylated arginine | 499.2251 | 499.2133 | 1.76 | 70.0654<br>114.1025<br>214.9622 | 5.70e3 |
| 3 | Glycosylated cysteine | 284.0804 | 284.0800 | 1.89 | 100.0216<br>122.0269<br>146.0267 | 3.93e6 |
| 4 | Glycosylated glutamine | 309.1298 | 309.1277 | 1.84 | 97.0285<br>130.0497<br>208.0599 | 3.71e5 |
| 5 | Glycosylated glutamic acid* | 310.1138 | 310.1120 | 1.83 | 97.0286<br>148.0603<br>226.0708 | 7.25e5 |
| 6 | Glycosylated isoleucine | 294.1552 | 294.1520 | 1.86 | 86.0966<br>132.1017<br>230.1382<br>258.1330 | 3.69e6 |
| 7 | Monoglycosylated lysine | 309.1661 | 309.1648 | 1.60 | 84.0810<br>128.0704<br>225.1227 | 4.50e6 |
| 8 | Diglycosylated lysine | 471.2189 | 471.2174 | 1.58 | 84.0810<br>212.1276<br>246.1330 | 1.29e5 |
| 9 | Glycosylated proline | 278.1240 | 278.1219 | 1.84 | 128.0705<br>242.1017<br>260.1122 | 2.29e5 |
| 10 | Glycosylated serine | 268.1032 | 268.1029 | 1.82 | 97.0286<br>130.0497<br>232.0811 | 4.88e5 |
| 11 | Glycosylated threonine | 282.1189 | 282.1171 | 1.81 | 97.0285<br>144.0653<br>246.0965 | 9.03e5 |
| 12 | Glycosylated valine | 280.1396 | 280.1389 | 1.80 | 216.1225<br>244.1173<br>262.1278 | 3.26e6 |
| 13 | Glycosylated alanine* | 252.1083 | 252.1071 | 1.82 | 102.0552<br>216.0866<br>234.0971 | 4.90e5 |
| 14 | Glycosylated asparagine | 295.1141 | 294.0034 | 1.83 | 87.0555<br>194.0444<br>211.0709 | 3.28e5 |
| 15 | Glycated aspartic acid | 296.0981 | 296.5928 | 2.05 | 85.0286<br>173.9607<br>260.0764 | 1.37e4 |
| 16 | Glycated glycine | 238.0927 | 239.1057 | 1.80 | 88.0395<br>97.0286<br>202.0706 | 1.06e5 |
| 17 | Glycated histidine | 317.1223 | 318.1276 | 1.58 | 95.0605<br>190.0970<br>238.1180 | 2.74e6 |
| 18 | Glycated leucine | 294.1552 | 294.0018 | 1.77 | 86.0967<br>132.1018<br>230.1383<br>258.1331 | 3.05e5 |
| 19 | Glycated methionine* | 312.1117 | 312.1121 | 1.84 | 88.0396<br>133.0317<br>276.0897 | 3.87e6 |
| 20 | Glycated phenylalanine | 328.1396 | 328.8529 | 1.89 | 186.9115<br>204.9221<br>288.8596 | 1.13e5 |
| 21 | Glycated tryptophan | 367.1505 | 367.1717 | 1.87 | 164.9297<br>186.9115<br>254.8987 | 6.50e3 |
| 22 | Glycated tyrosine | 344.1345 | 345.1556 | 1.87 | 165.0541<br>281.8988<br>308.1132 | 8.11e3 |
| 23 | Glycated creatine | 294.1301 | 294.1369 | 1.61 | 90.9771<br>158.9640<br>206.9552 | 5.28e5 |
| 24 | Monoglycated urea | 223.0930 | 223.0938 | 1.71 | 104.9925<br>116.9925<br>135.0029 | 5.79e6 |
| 25 | Diglycated urea | 385.1458 | 385.1205 | 1.80 | 203.0526 | 6.09e5 |

\*Glycated amines synthesized on 7<sup>th</sup> day

**Table 2:** HRMS analysis of glycated amines detected in the diabetic plasma.

| S. No. | Glycated amines | Monoisotopic mass (m/z) | Observed mass (m/z) | RT (min) | Fragment ions matching with *in vitro* synthesized |
| --- | --- | --- | --- | --- | --- |
| 1 | Glycated arginine | 337.1723 | 337.1719 | 1.61 | 60.0561<br>70.0656<br>114.0915<br>173.1397<br>217.1292<br>257.1604 |
| 2 | Glycated lysine | 309.1661 | 309.1656 | 1.67 | 84.0810<br>128.0705<br>225.1230 |
| 3 | Glycated isoleucine | 294.1552 | 294.2134 | 2.06 | 86.0967<br>132.1018<br>230.1384<br>258.1332 |
| 4 | Glycated urea | 223.0930 | 223.0937 | 1.80 | 104.9924<br>116.9923<br>135.0027 |
| 5 | Diglycated urea | 385.1458 | 385.1518 | 1.90 | 203.0523 |

### 2.6. Quantification of glycated urea by PRM

Targeted quantification of glycated urea was performed by PRM [9]. The mass spectrometric raw data were processed and analyzed using Skyline (version 21.2.0.568) with the help of the transitions of metabolites of interest.

2µl of ^13^C6-glycated urea (0.4mM) was used as the internal standard for normalization. Relative quantification of glycated urea was performed by the following equation.

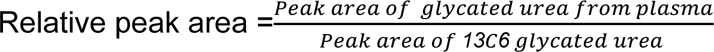

^12^C6-Glycated urea and ^13^C6-Glycated urea were synthesized by incubating 500 µl of 100 mM urea with 500 µl of 500 mM of ^12^C6-glucose and ^13^C6-glucose in 50 mM phosphate buffer (pH 7.4) for 72 h at 37℃, respectively. A tandem mass spectrometry (MS/MS) was performed for both ^13^C6 and ^12^C6-glycated urea to characterize the fragmentation pattern.

### 2.7. Correlation of glycated urea with various clinical parameters

Correlation analysis of plasma glycated urea was carried out with all the recorded clinical parameters like fasting and postprandial blood glucose levels, HbA1c, microalbuminuria, and serum creatinine using Pearson’s correlation method. Here, the data of different subject groups (H, DM, DN) were analyzed using GraphPad Prism 8.0.2 software. In the present study, p-values less than 0.05 were considered statistically significant [9].

## 3. Results

### 3.1. *In vitro* synthesis of glycated amino acids

The *in vitro* reaction between glucose and various amino acids for three days led to the synthesis of 10 glycated amino acids that are listed in Table 1. Tandem mass spectrometric analysis of these glycated amino acids provided information about their fragments, which will be useful for identifying and quantifying glycated amino acids from complex matrices like plasma. Detailed information on the precursor and fragments of glycated amino acids is provided in Table 1. Amongst these glycated amino acids, lysine, cysteine, isoleucine, and valine showed higher intensity (Table 1, Fig. S1), suggesting these are more prone to undergo modification by glucose. Alternatively, these glycated amino acids may have higher ionization efficiency. Also, glutamic acid, alanine, and methionine were found to be glycated on the 7th day (Table 1), suggesting these amino acids are relatively less susceptible to glycation.

In addition to amino acids, other small molecule amines, such as urea and creatine were also found to be glycated *in vitro* (Table 1). Urea was detected in both monoglycated and diglycated forms (Fig. 2). creatine, a precursor of creatinine, and urea were considered for the *in vitro* glycation reaction, as these are markers of kidney dysfunction and diabetic nephropathy [6, 10].

### 3.2. Identification of glycated amines in the plasma of diabetic patients

The presence of glycated amines in the plasma was confirmed by matching their precursor and fragment ions with the *in vitro* synthesized glycated amines. By this approach, the presence of glycated forms of lysine, arginine, leucine/isoleucine was detected in the diabetic plasma. Glycated lysine (m/z = 309.1661) from the plasma showed the following matching fragments (m/z = 84.08, 128.07, 225.12) with the in vitro synthesized glycated lysine. Similarly, glycated arginine (m/z = 337.1723) and glycated leucine/isoleucine (m/z = 294.1552) showed matching fragments (m/z = 60.06, 70.07, 114.09, 173.14, 217.13, 257.16) and (m/z = 86.1, 132.10, 230.14, 258.13) with their *in vitro* forms, respectively (Table 2, Fig. 1 and Fig S2). Apart from these three amino acids, the glycated forms of other amino acids were not detected in the plasma. Diabetic plasma was used to detect glycated amines with the assumption that their abundance would be higher in the hyperglycemic conditions.

**Fig. 1.**
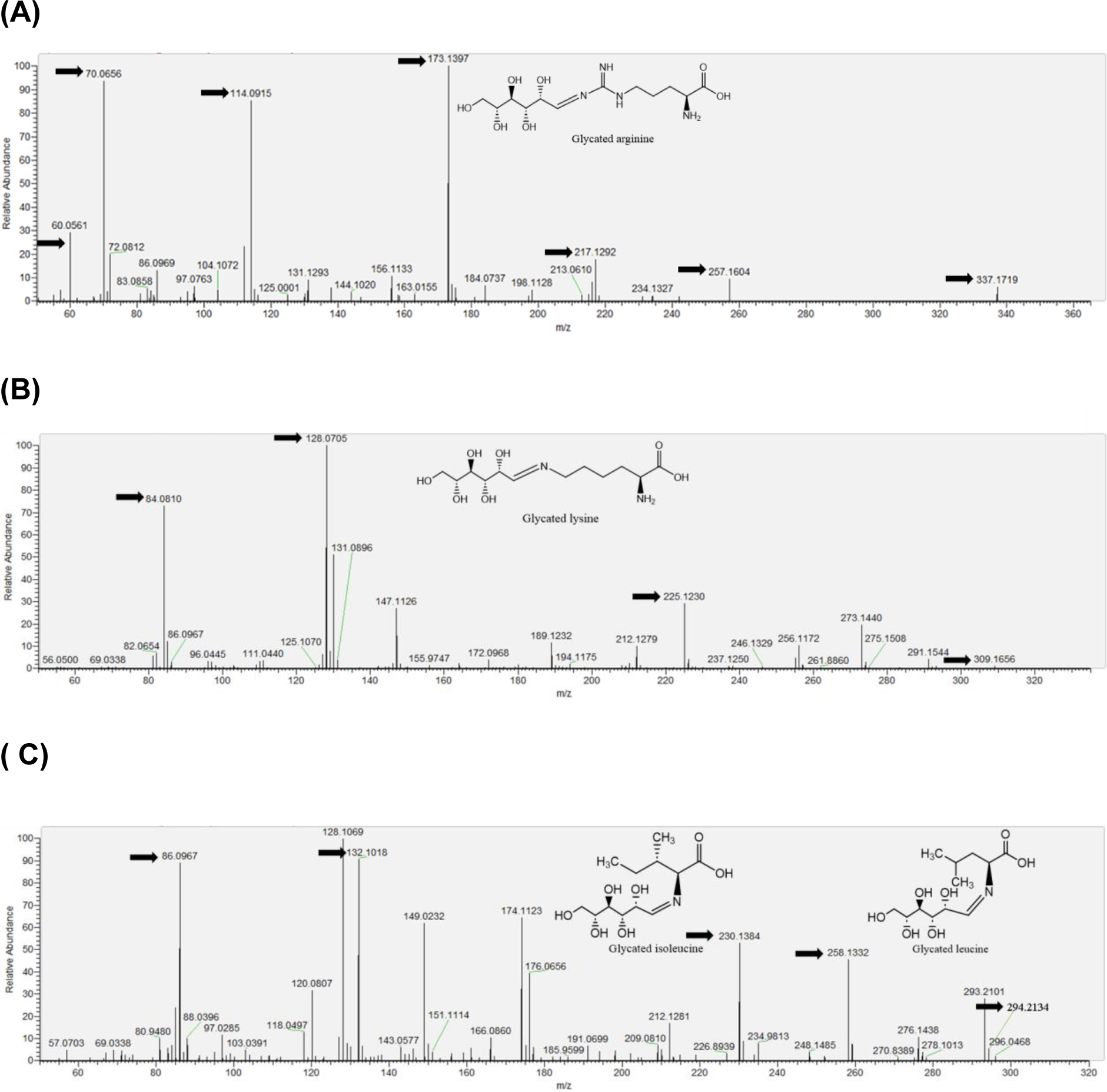
Mass spectrum of glycated amino acids derived from diabetic plasma samples (Predicted structures are shown in the inset). Matching fragments are indicated with the arrows. (A) Detected glycated arginine from diabetic plasma samples (B) Detected glycated lysine from diabetic plasma samples (C) Detected glycated isoleucine/leucine from diabetic plasma samples

**Fig. 2.**
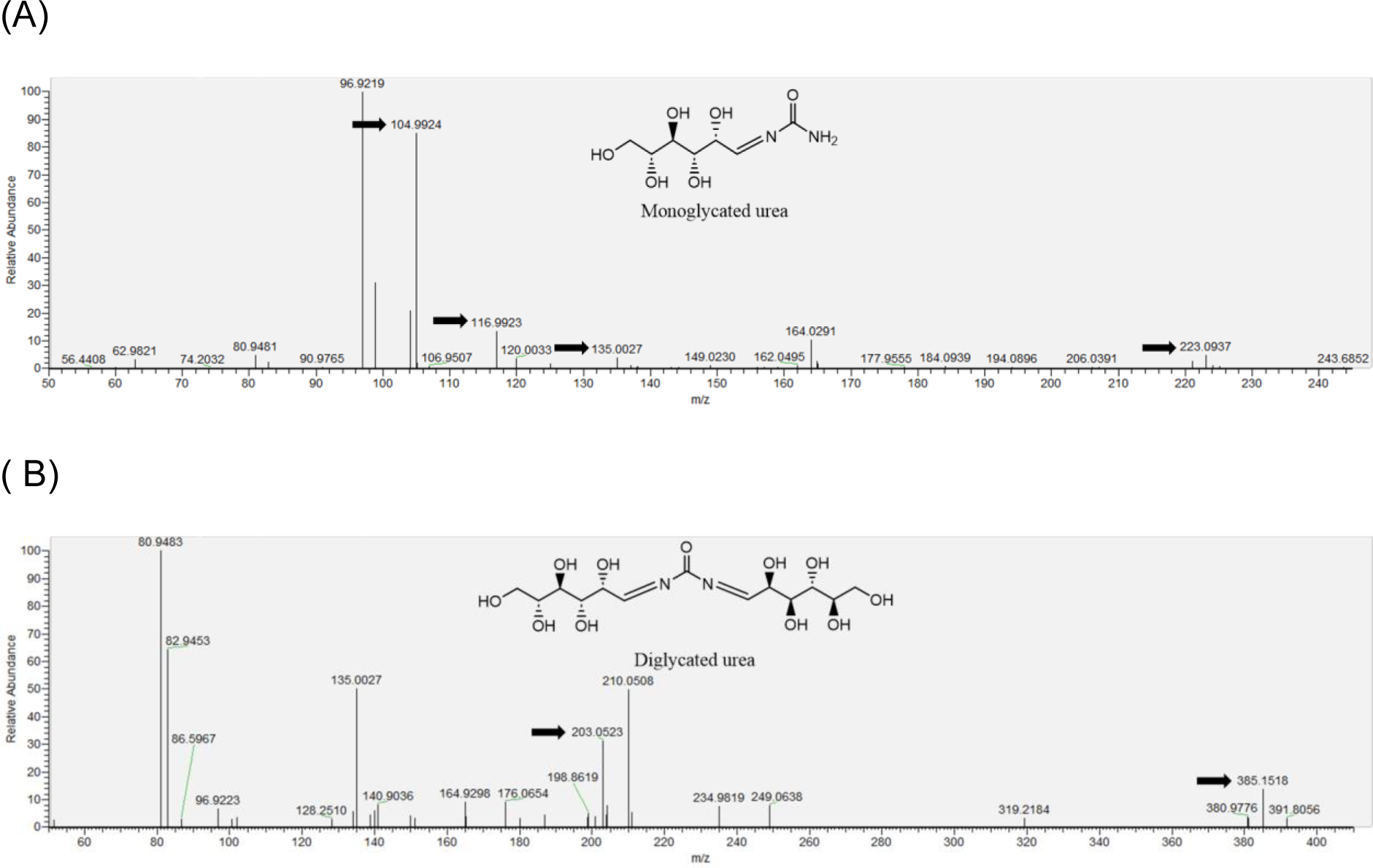
Mass spectrum of (A) monoglycated urea (MGU) and (B) diglycated urea (DGU) from diabetic plasma. Matching fragments are indicated with the arrows.

Besides glycated amino acids, glycated urea was detected in the diabetic plasma. The precursor (m/z= 223.0930) and fragment masses (m/z = 104.99, 116.99, 135.00) for monoglycated urea (MGU) from the diabetic plasma matched with the *in vitro* synthesized MGU (Table 2, Fig. 2 and Fig S3A). Similarly, m/z = 385.1458 corresponding to DGU showed the following fragment matches (m/z = 203.05) with the synthetic DGU. (Table 2, Fig. 2 and Fig S3B). However, we could not detect the glycated forms of creatine in the plasma.

### 3.3. Quantification of MGU and DGU by PRM

Blood urea nitrogen is one of the diagnostic markers for kidney dysfunction and diabetic nephropathy [9]. Currently, urea is quantified enzymatically by using urease, which may not detect glycated urea, thus affecting the accuracy of diagnosis. Therefore, in this study, MGU and DGU were quantified by PRM in healthy (n=26), diabetic (n=26), and diabetic nephropathy (n=23) subjects. The mean peak area of MGU (8.8 x 105) was more abundant than the peak area of DGU (1.7 x 105) in all the subjects (Table S1). For normalization, ^13^C6-MGU and ^13^C6-DGU were synthesized using ^13^C6- glucose and urea, and their mass spectral fragmentation was compared to ^12^C6-MGU and ^12^C6-DGU. The MS/MS spectra of ^12^C6-MGU and ^12^C6-DGU showed a similar fragmentation pattern to that of ^13^C6-MGU and ^13^C6-DGU, respectively (Fig S4A and Fig S4B). Therefore, the peak area of *in vitro* synthesized ^13^C6-MGU and ^13^C6-DGU, was used for normalization.

A representative Skyline generated peak area for MGU and DGU is depicted in Fig 4A and Fig 4C, respectively. The normalized AUC of MGU (mean ± SD) in healthy, diabetes and DN was 1.01 ± 0.29, 1.62 ± 0.65, and 1.54 ± 0.76, respectively (Table S1). The normalized mean MGU area was significantly higher in DM and DN compared to healthy subjects. However, the MGU levels were not significantly different between DM and DN (Fig 4B). The DGU also showed a similar trend in H, DM, and DN. The normalized AUC of DGU was significantly higher in DM and DN (Fig. 4D, Table S1). However, the values of DGU appear to be higher than that of MGU, although MGU was more abundant. This is due to the lower detection of ^13^C6-DGU in the mass spectrometric analysis and its use in normalization.

**Fig 3.**
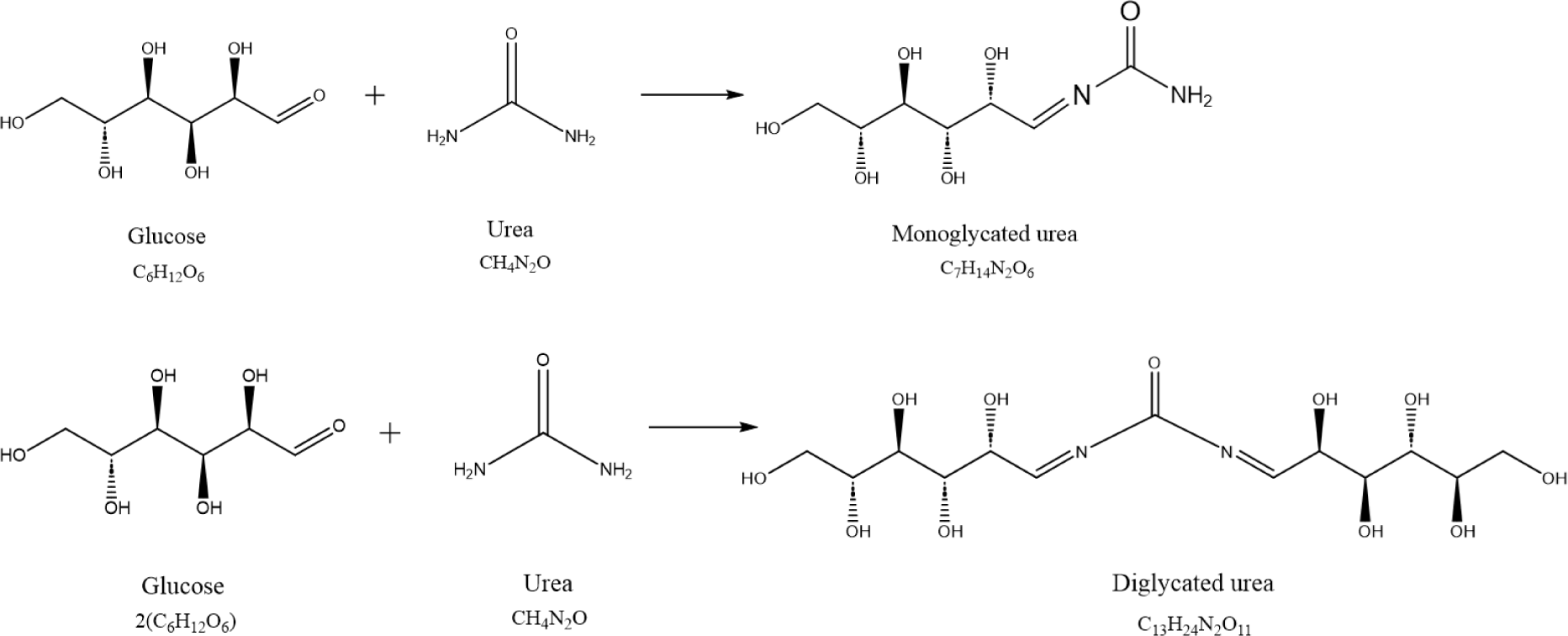
Proposed glycation reaction for (A) monoglycated urea (MGU) and (B) diglycated urea (DGU) synthesis

**Fig. 4.**
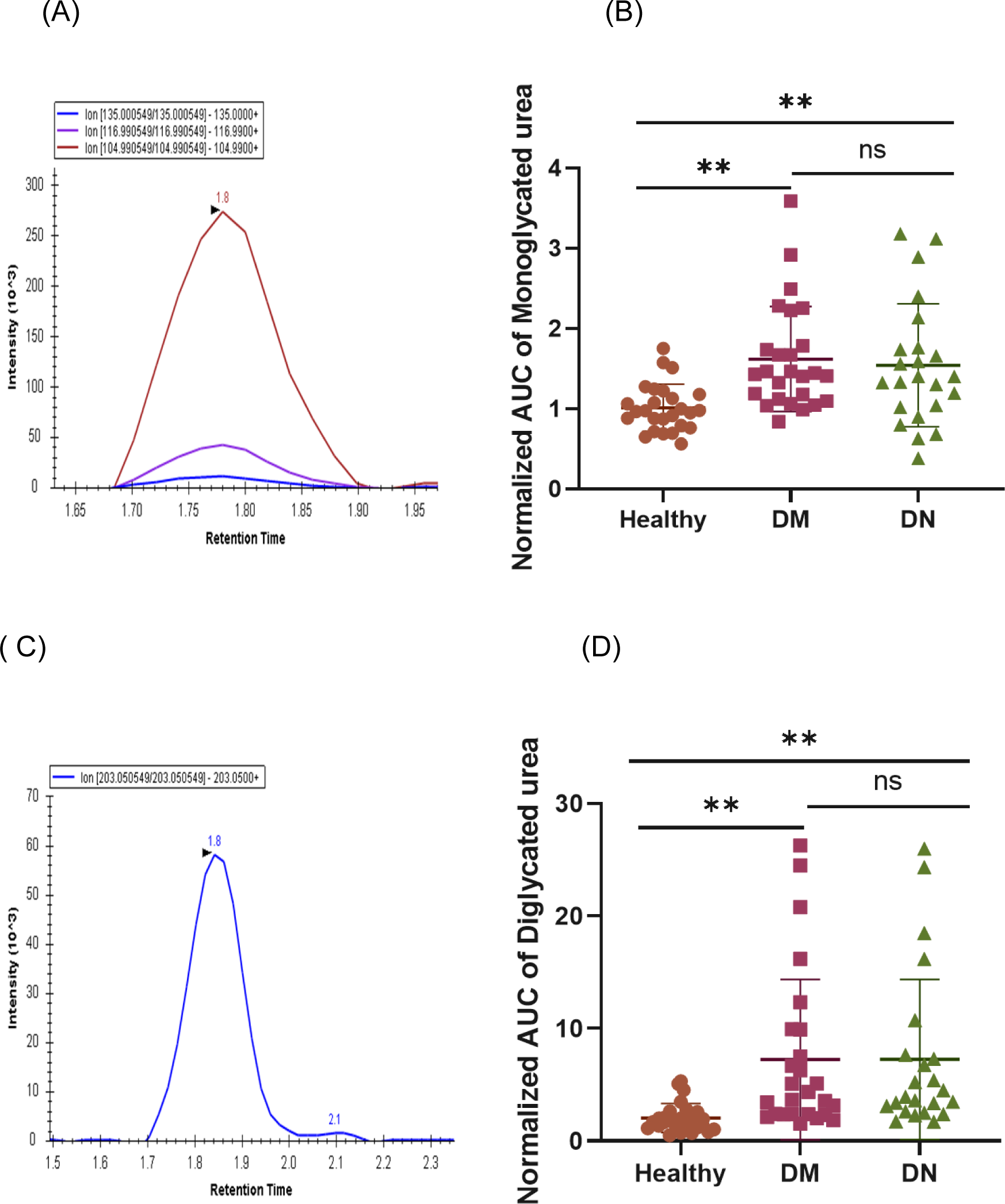
Representative chromatogram of (A) monoglycated urea (MGU) obtained from Skyline analyses (B) Normalized AUC of MGU in healthy, diabetic and diabetic nephropathy plasma samples (C) Diglycated urea (DGU) obtained from Skyline analyses (D) Normalized AUC of DGU in healthy, diabetic and diabetic nephropathy plasma samples

Furthermore, the abundance of MGU and DGU was correlated with the various clinical parameters. Both MGU and DGU showed significant correlations with fasting blood glucose, postprandial glucose, and HbA1c. MGU showed a significant correlation with fasting blood sugar level (r= 0.78***), postprandial blood sugar level (r= 0.64***), and HbA1c (0.62***) (Table 3). DGU also showed a significant correlation with fasting blood sugar level (r= 0.74***), postprandial blood sugar level (r= 0.62***), and HbA1c (r= 0.58***) (Table 3). However, both did not show a strong correlation with the known markers of kidney dysfunctions, such as microalbumin and serum creatinine.

**Table 3:** Correlation of glycated urea with clinical parameters.

| Clinical parameters | MGU |  | DGU |  |
| --- | --- | --- | --- | --- |
|  | *r* | *p* value | *r* | *p* value |
| HbA1c | 0.62 | <0.0001 | 0.58 | <0.0001 |
| FBSL | 0.78 | <0.0001 | 0.74 | <0.0001 |
| PP BSL | 0.64 | <0.0001 | 0.62 | <0.0001 |
| Serum creatinine | 0.004 | 0.97 | -0.029 | 0.8 |
| Microalbumin | 0.135 | 0.29 | 0.184 | 0.14 |

†FBSL: Fasting blood sugar level, PPBSL: Postprandial blood sugar level

## 4. Discussion

An enormous amount of research exists concerning the glycation of proteins in diabetes. However, the information on glycation of free small-molecule amines is very scanty. There are a few reports of small-molecule glycation in yeast [8]. However, to the best of our knowledge, we have not come across any information regarding the glycation of small molecule amines in humans. In this quest, free glycated amino acids and glycated urea were discovered from the diabetic plasma in this study. Glycated amino acids and glycated urea were synthesized *in vitro,* and a mass spectral library comprising precursor and fragment masses was developed. This information was used to identify free glycated amino acids and glycated urea from the diabetic plasma. For the first time, we report the identification of glycated lysine, glycated arginine, and glycated leucine/isoleucine from the diabetic plasma. The presence of these glycated amino acids in plasma was validated by comparing their precursor and fragment masses with the synthetic glycated amino acids. The discovery of free glycated amino acids has great physiological significance. Glycation of amino acids may create a condition akin to amino acid deficiency, as the glycated amino acids may not be suitable to carry out the functions of amino acids. Amino acids are essential for protein synthesis, and their deficiency may affect the protein synthesis. In diabetic conditions, there is an overall decline in protein content and muscle mass [12]. This decrease is attributed to increased protein catabolism due to insulin deficiency. However, based on this study, we propose that the glycation of amino acids leads to the deprivation of amino acids and affects protein synthesis. Besides their role in protein synthesis, amino acids are involved in cell signaling and metabolic regulation. Amino acid abundance is known to activate mTOR1 pathway, which in turn, through a cascade of signaling, induces insulin secretion. In contrast, the deficiency of amino acids activates GCN2, which inactivates mTOR1 pathway, thereby affecting insulin secretion [13, 14]. A decrease in insulin secretion may lead to hyperglycemic conditions and glycation of amino acids, thereby forming a vicious cycle of amino acid deficiency and activation of GCN2 pathway.

Furthermore, glycation of these individual amino acids, lysine, arginine, leucine/isoleucine in hyperglycemic conditions may create their deficiency, consequently, their deficiency symptoms may be observed in diabetic conditions. Lysine is an essential amino acid known for positive nitrogen balance in the human body through its abundance in different human proteins [15]. Lysine deficiency results in a decrease in body weight gain [16]. A lysine-deficient diet reduces growth and bone metabolism, which impacts calcium intestinal absorption and collagen synthesis [17]. Lysine has antihyperglycemic effects and can improve diabetic conditions in humans and rats by reducing blood sugar and non-enzymatic glycation. Two studies revealed that lysine postponed cataracts, as a complication of DM, by lowering blood glucose [18,19]. Besides its actions as a chemical chaperone, L-lysine can reduce inflammation by lowering the levels of interleukin 4 in the kidney and interleukin 10 in the liver. Some studies showed that L-lysine limitation activated NF-*κ*B and ERK1/2 signals and caused inflammation [20]. Glycation of lysine may affect its useful functions in diabetes. Arginine, a functional amino acid, the precursor of nitric oxide, plays a crucial role in the maintenance, reproduction, growth, anti-aging, and immunity of animals. It is synthesized from glutamate, glutamine, and proline via citrulline, whereas it is degraded to nitric oxide, ornithine, urea, polyamines, proline, glutamate, creatine, and agmatine through various pathways. In fact, a large amount of arginine is catabolized to creatine, which is a potent antioxidant and improves glucose tolerance [20]. Hyperglycemic conditions in ketosis-prone diabetes have been shown to be associated with decreased arginine availability and decreased insulin secretion, whereas exogenous arginine supplementation restored insulin secretion [21]. Other studies have also reported the induction of insulin secretion by arginine [22]. Numerous studies conducted on human and animal models have found that arginine supplementation improves insulin sensitivity in diabetes [23].

Isoleucine/ leucine are branched-chain amino acids (BCAA), which are essential for protein synthesis, energy production, performing signaling functions mainly through mTOR pathway, and also act as the source of nitrogen for the synthesis of glutamate, glutamine, alanine, and aspartate [24]. The concentration of BCAA tends to decrease in hyperammonemia states like liver cirrhosis and urea cycle disorders [24]. Although in diabetes, the levels of BCAAs are higher, however, they can undergo glycation and, as a result, may not be available for ammonia detoxification to glutamine (GLN). Leucine and isoleucine are also crucial for stimulating β-cell electrical activity, which is essential for upregulating glucose transporters and insulin secretion, and glycation may affect this process.

Apart from affecting key physiological functions due to glycation, these glycated amino acids can potentially serve as a marker for diabetes and its complications, which requires it to be studied in detail. Diabetic nephropathy is one of the major diabetic complications. Currently, it is diagnosed by microalbuminuria, serum creatinine clearance, and blood urea nitrogen [10]. Creatinine is formed from dehydration and dephosphorylation of creatine[26]. Creatine is a small molecule amine with a primary amino group capable of undergoing glycation, consequently, may lead to lesser production of creatinine due to the non-availability of unmodified creatine. This may underestimate creatinine accumulation in the serum despite the worsening of kidney function in diabetes. However, in this study, we could not detect glycated creatine in the diabetic plasma. Similarly, blood urea nitrogen is also used to diagnose diabetic nephropathy [10]. During detoxification of ammonia, arginine is converted to ornithine and urea. Urea is a small molecule amine with two primary amino groups and is susceptible to glycation modification in hyperglycemic conditions. In this study, for the first time, we report the discovery of the formation of glycated urea, which was detected in monoglycated and diglycated forms in the diabetic plasma. The abundance of both monoglycated urea (MGU) and diglycated urea (DGU) was found to be higher in the plasma of diabetes and diabetic nephropathy. Both MGU and DGU were significantly correlated with fasting blood glucose, postprandial glucose, and HbA1c. Although neither MGU nor DGU showed a significant correlation with the markers of diabetic nephropathy, their formation may lead to an underestimation of urea in diabetic nephropathy. Therefore, quantification of glycated urea could be useful to detect early kidney disease. The usefulness of quantification of glycated urea needs to be studied in a larger cohort. Also, it is important to study the physiological role of glycated urea, including MGU and DGU accumulation in diabetes. Additionally, it would be interesting to quantify glycated urea in Uremic conditions. Apart from these, the other biogenic amines such as serotonin, tryptamine, ornithine, polyamines like spermine and spermidine, GABA etc., are also susceptible to *in vivo* glycation in hyperglycemic conditions. It would be interesting to study the effect of glycation on these molecules in diabetic conditions.

## Conclusions

In this study, for the first time, we report the discovery of glycation of free lysine, arginine, leucine/isoleucine from the diabetic plasma. This has great physiological significance as glycation of these amino acids may affect their vital functions such as protein synthesis, cell signaling and insulin secretion. Also, the glycated amino acids could be used as potential markers for the diagnosis and management of diabetes and its complications. Diabetic nephropathy is one such complication, where amines such as creatinine and urea accumulate in the plasma. For the first time, we report the detection of glycated urea in diabetic plasma. Further, we quantified MGU and DGU by targeted mass spectrometric approach in the plasma of healthy, diabetes, and diabetic nephropathy subjects. Both MGU and DGU show a strong correlation with clinical parameters such as blood glucose and HbA1c. Given that urea gets converted to glycated urea in hyperglycemic conditions while quantifying urea for kidney diseases, glycated urea needs to be accounted for.

## Acknowledgement

This work is supported by CSIR project HCP47, Chellaram Diabetes Research Center grant CDRC202111009. MJK and SD thank the DBT (Govt. of India) BioCARE program (Project code:BT/PR50872/BIC/101/1294/2023).

## Author Contribution

**Rashdajabeen Q Shaikh:** Data curation, Formal analysis, Investigation, Methodology, Writing - review & editing; **Sancharini Das:** Data curation, Formal analysis, Investigation, Methodology, Writing - review & editing; **Arvindkumar Chaurasiya:** Methodology, Writing - review & editing, Validation; Visualization; **Ashtamy MG**: Formal analysis, Validation, Writing - review & editing; **Amreen B Sheikh**: Writing - review & editing, Validation; **Moneesha Fernandes**: Writing - review & editing, Validation; **Shalbha Tiwari**: Investigation, Resources Writing - review & editing; **Unnikrishnan AG**: Investigation, Resources, Writing - review & editing; **Mahesh J Kulkarni**: Conceptualization; Investigation; Methodology; Project administration, Supervision; Writing - original draft.

## Supplementary information

List of supplementary figures

Fig S1. MS/MS spectra of *in vitro* synthesized glycated amino acids.

Fig S2. MS/MS spectra of glycated amino acids detected in diabetic plasma and their corresponding *in vitro* synthesized glycated amino acids with the matching fragments.

(A) *In vitro* synthesized glycated arginine (B) glycated arginine from diabetic plasma (C) *in vitro* synthesized glycated lysine (D) glycated lysine from diabetic plasma) (E) *in vitro* synthesized glycated isoleucine (F) glycated isoleucine/leucine from diabetic plasma samples

Fig S3. MS/MS spectra of glycated MGU and DGU detected in diabetic plasma and their corresponding *in vitro* synthesized MGU and DGU with the matching fragments.

(A) *In vitro* synthesized MGU (B) MGU from diabetic plasma (C) *in vitro* synthesized DGU (D) DGU from diabetic plasma

Fig S4. (A) MS/MS spectra of ^12^C6-MGU and ^13^C6-MGU (B) MS/MS spectra of ^12^C6- DGU and ^13^C6-DGU

## List of supplementary Tables

Table S1. Detailed information of cumulative mean and normalized mean in all subjects

