## Supplementary material for "Discovery of free glycated amines and glycated urea as potential markers of diabetic nephropathy": suppl information

### Slide 1
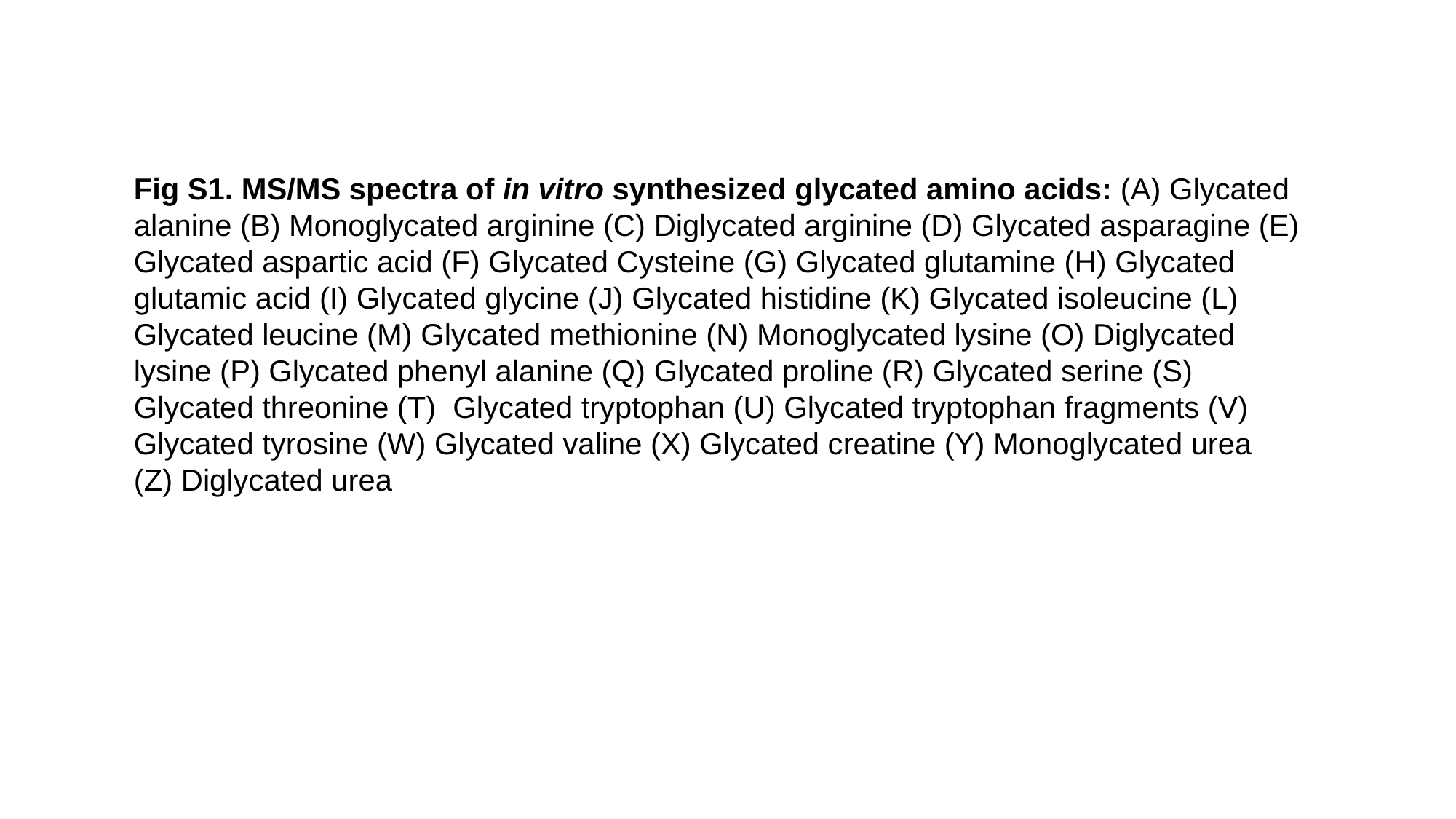

Fig S1. MS/MS spectra of in vitro synthesized glycated amino acids: (A) Glycated alanine (B) Monoglycated arginine (C) Diglycated arginine (D) Glycated asparagine (E) Glycated aspartic acid (F) Glycated Cysteine (G) Glycated glutamine (H) Glycated glutamic acid (I) Glycated glycine (J) Glycated histidine (K) Glycated isoleucine (L) Glycated leucine (M) Glycated methionine (N) Monoglycated lysine (O) Diglycated lysine (P) Glycated phenyl alanine (Q) Glycated proline (R) Glycated serine (S) Glycated threonine (T) Glycated tryptophan (U) Glycated tryptophan fragments (V) Glycated tyrosine (W) Glycated valine (X) Glycated creatine (Y) Monoglycated urea
(Z) Diglycated urea

### Slide 2
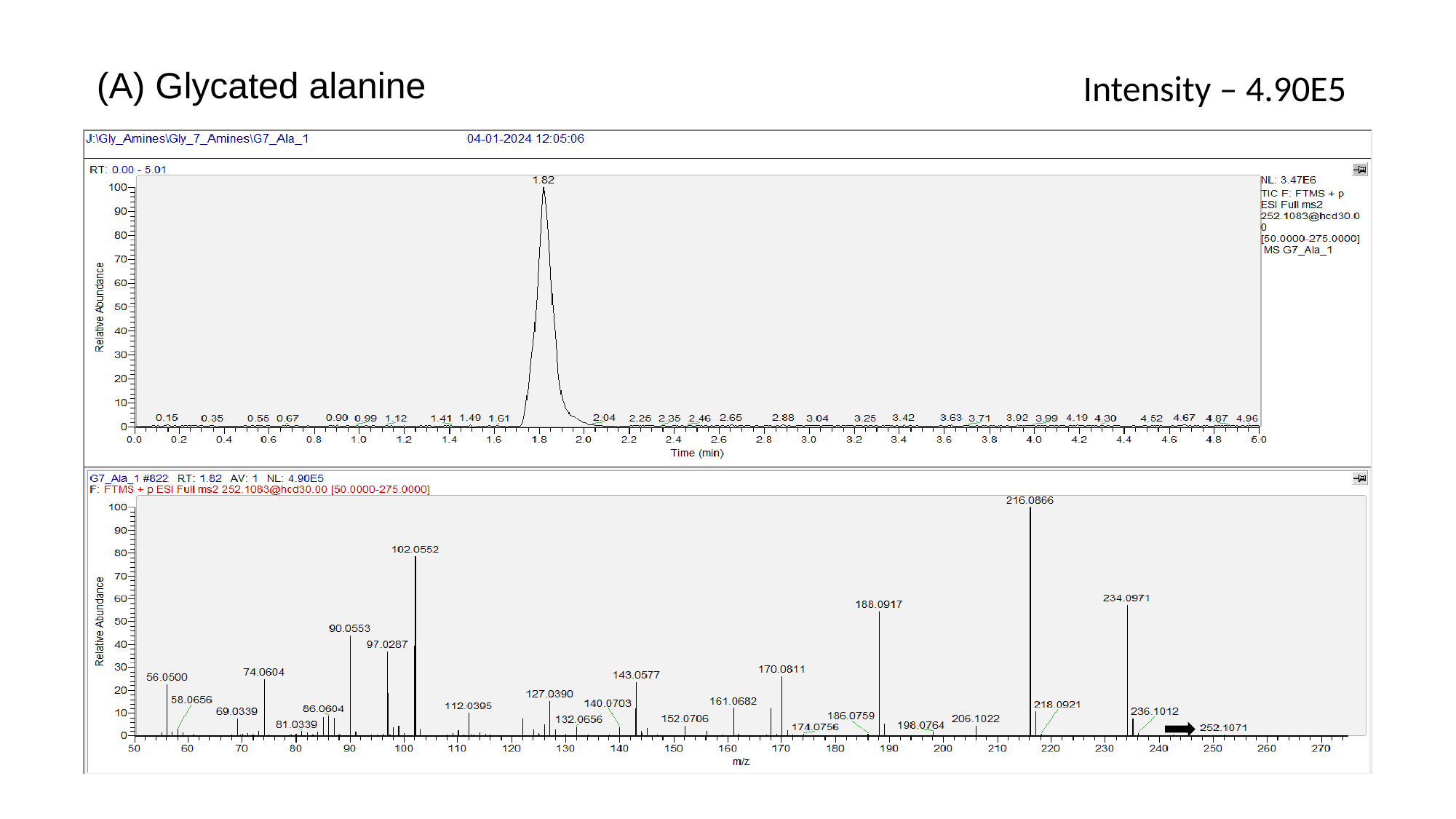

(A) Glycated alanine
Intensity – 4.90E5

### Slide 3
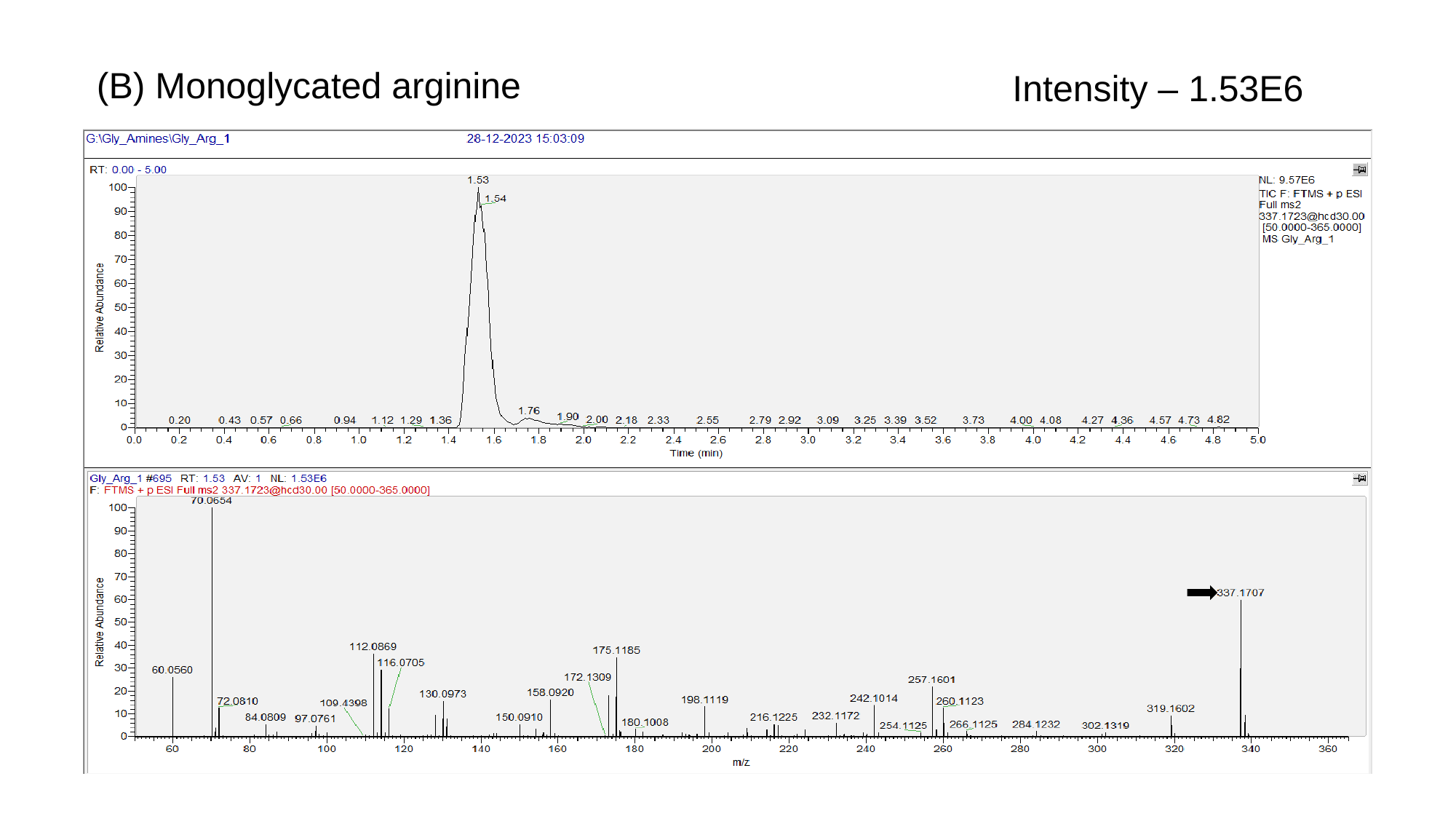

(B) Monoglycated arginine
Intensity – 1.53E6

### Slide 4
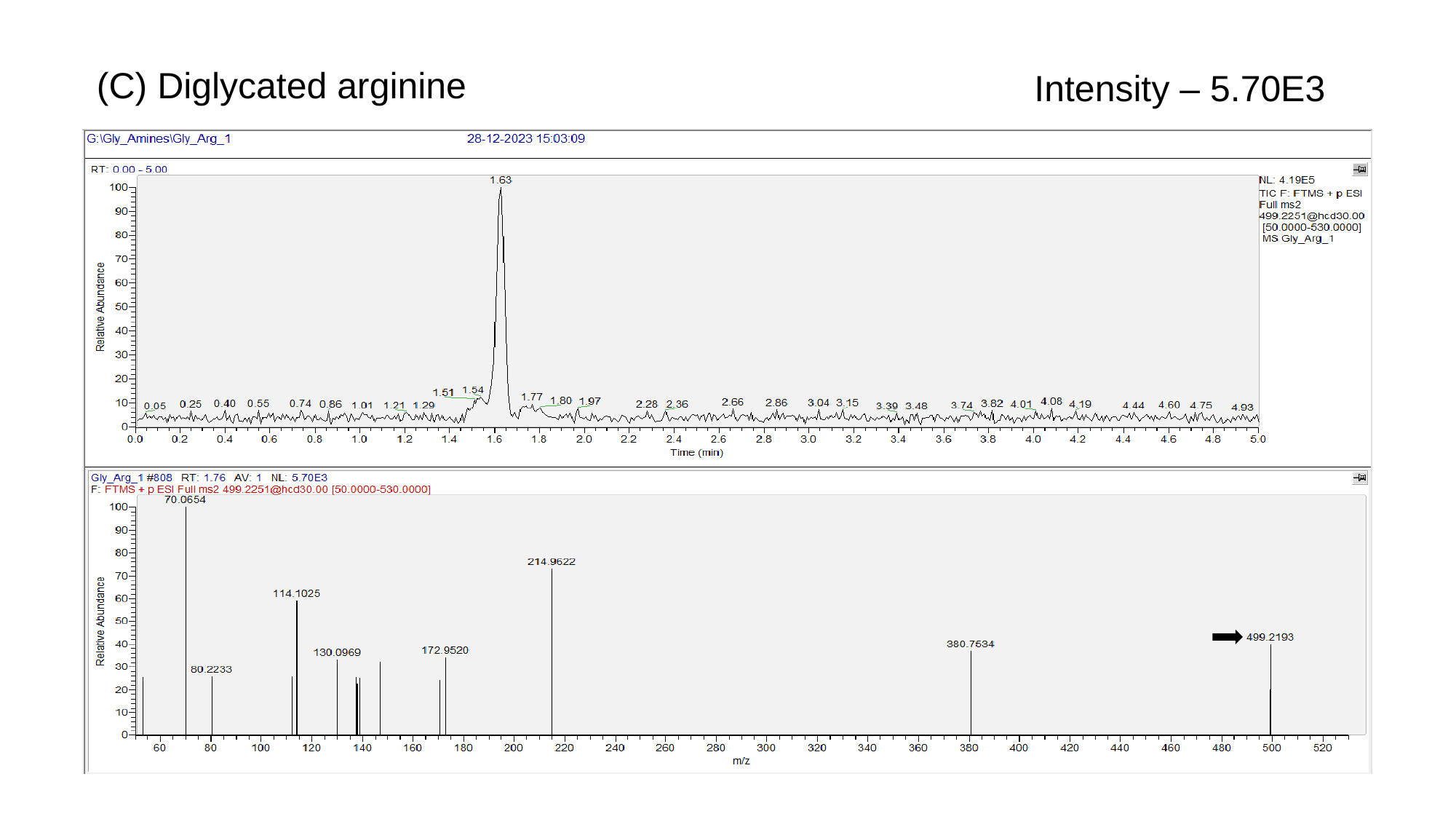

(C) Diglycated arginine
Intensity – 5.70E3

### Slide 5
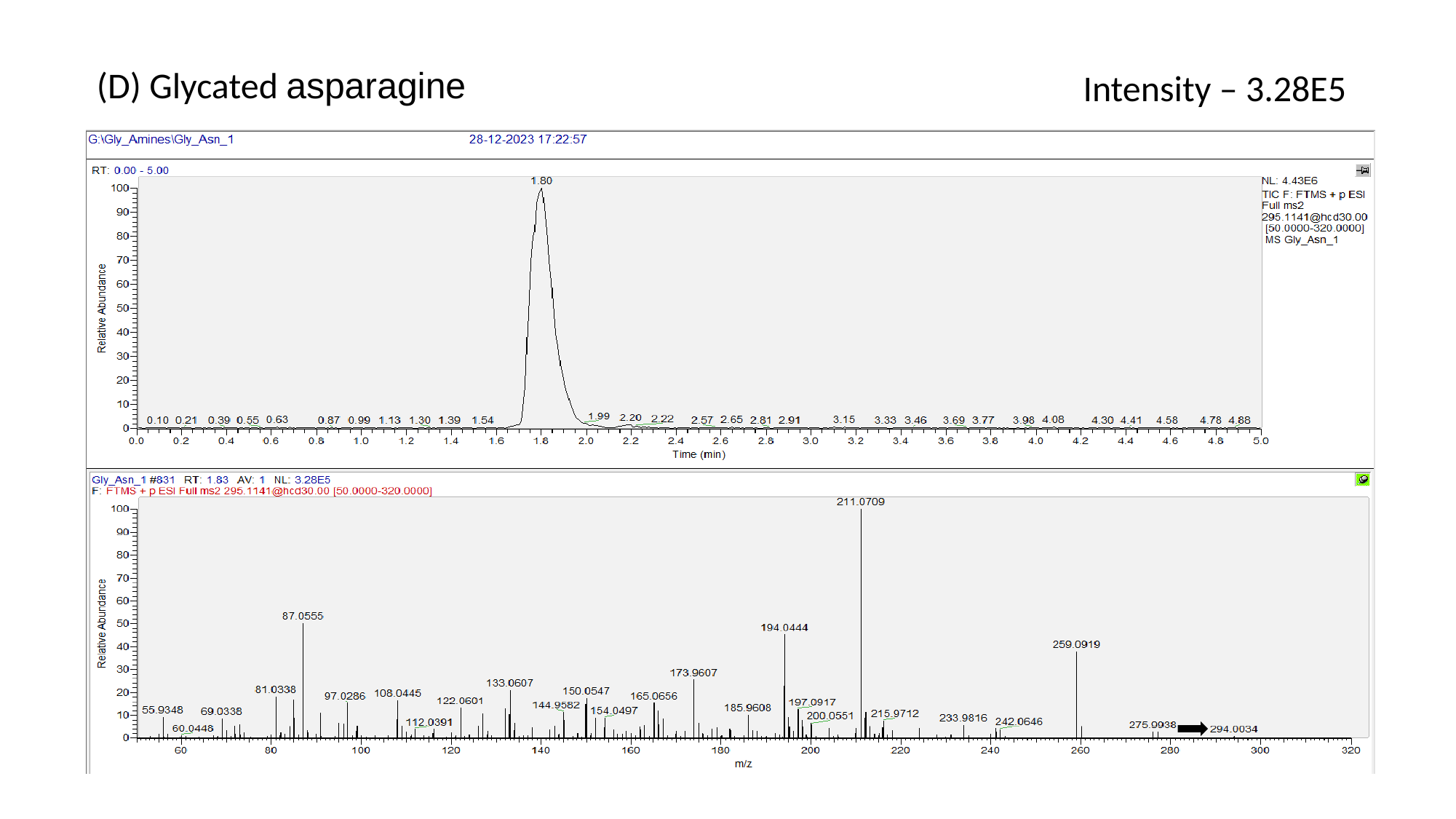

(D) Glycated asparagine
Intensity – 3.28E5

### Slide 6
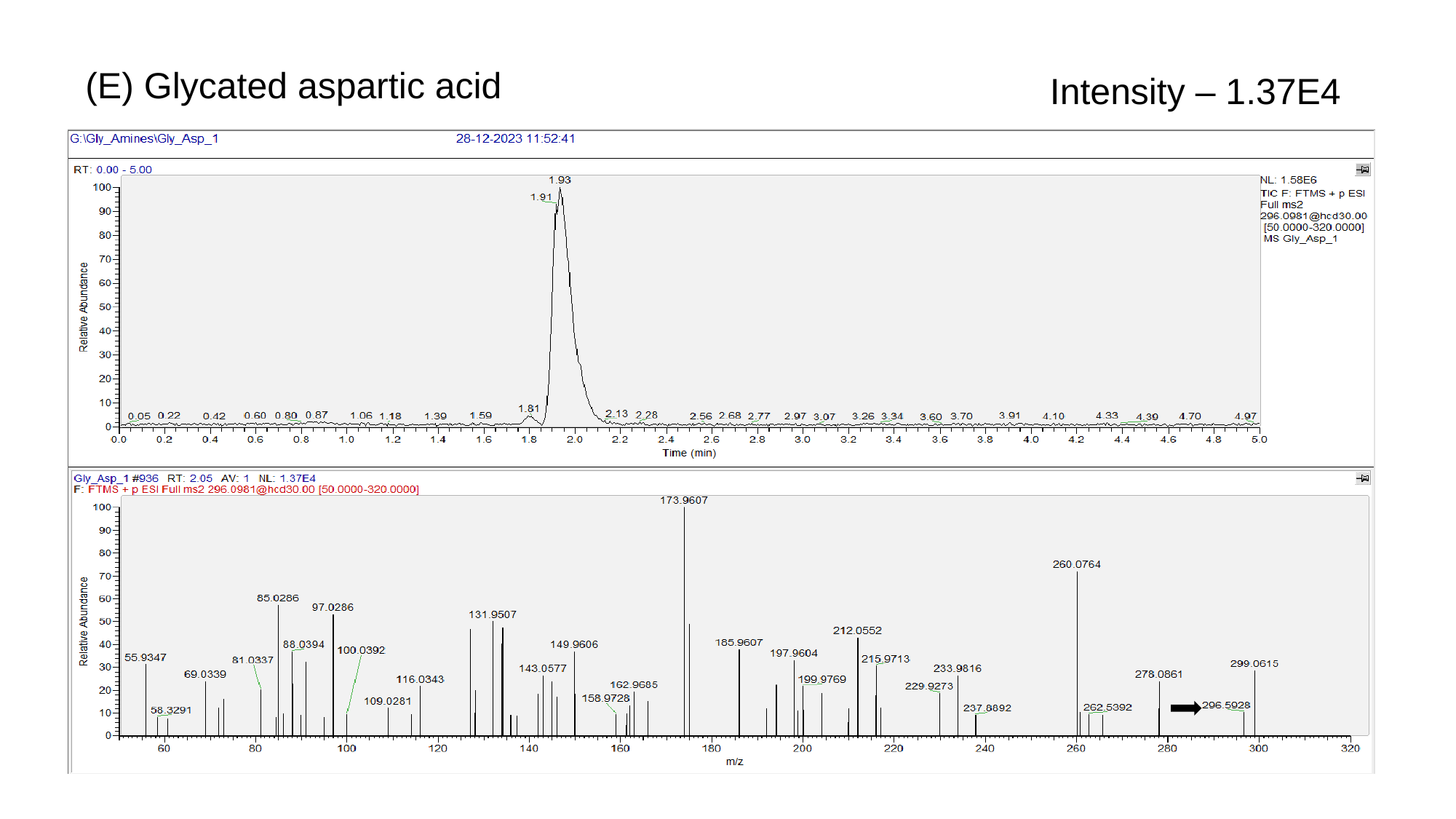

(E) Glycated aspartic acid
Intensity – 1.37E4

### Slide 7
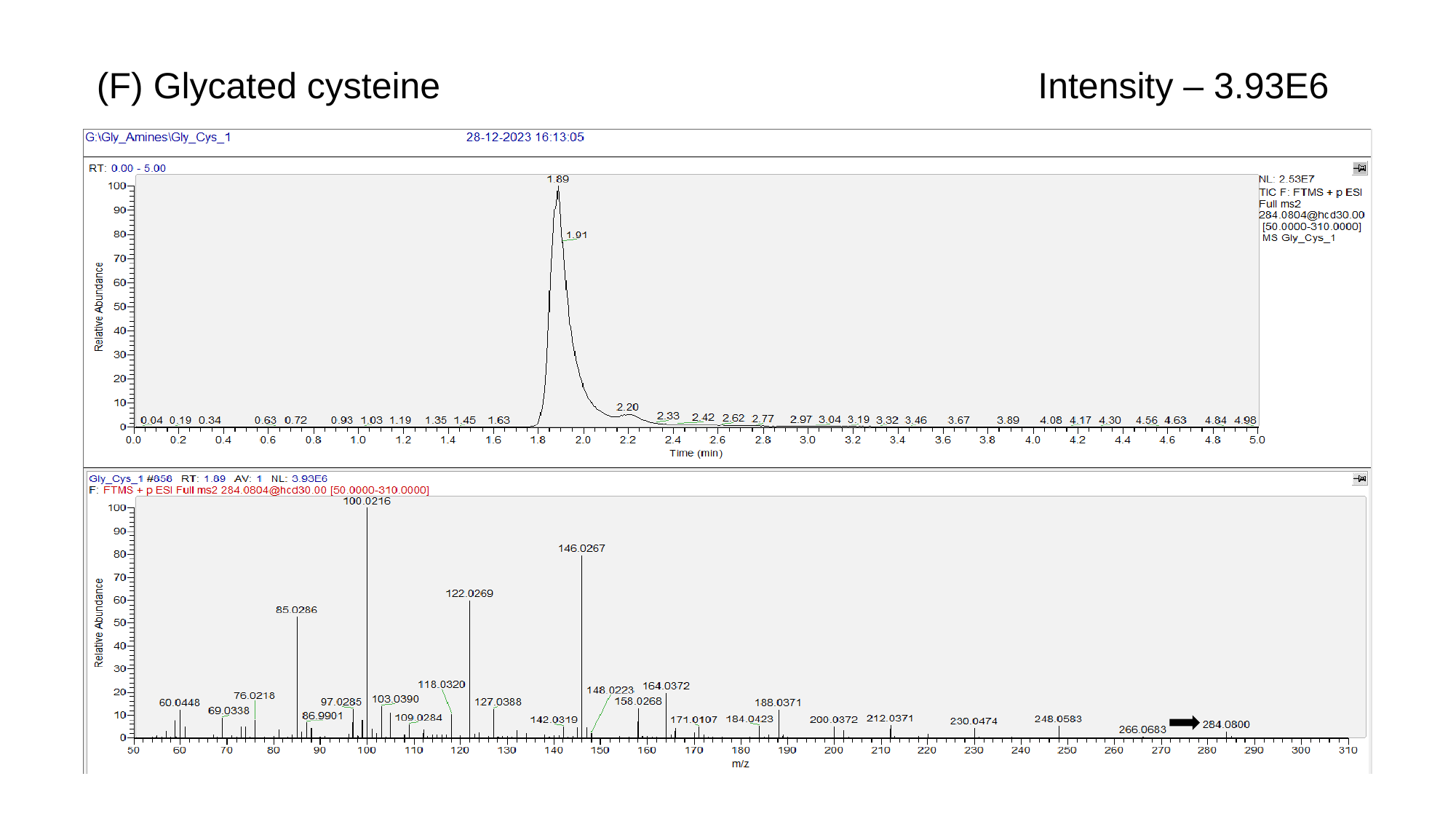

(F) Glycated cysteine
Intensity – 3.93E6

### Slide 8
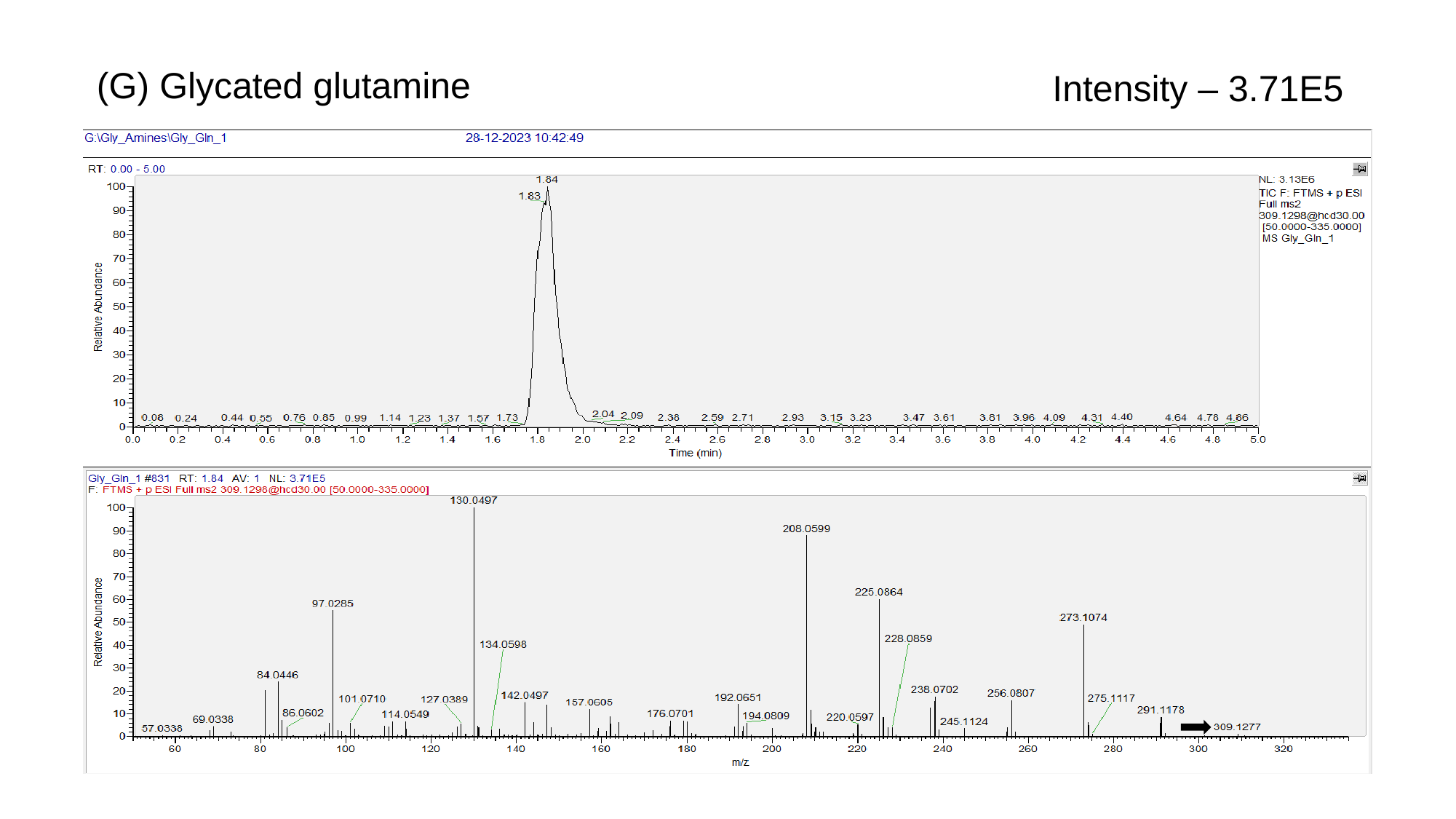

(G) Glycated glutamine
Intensity – 3.71E5

### Slide 9
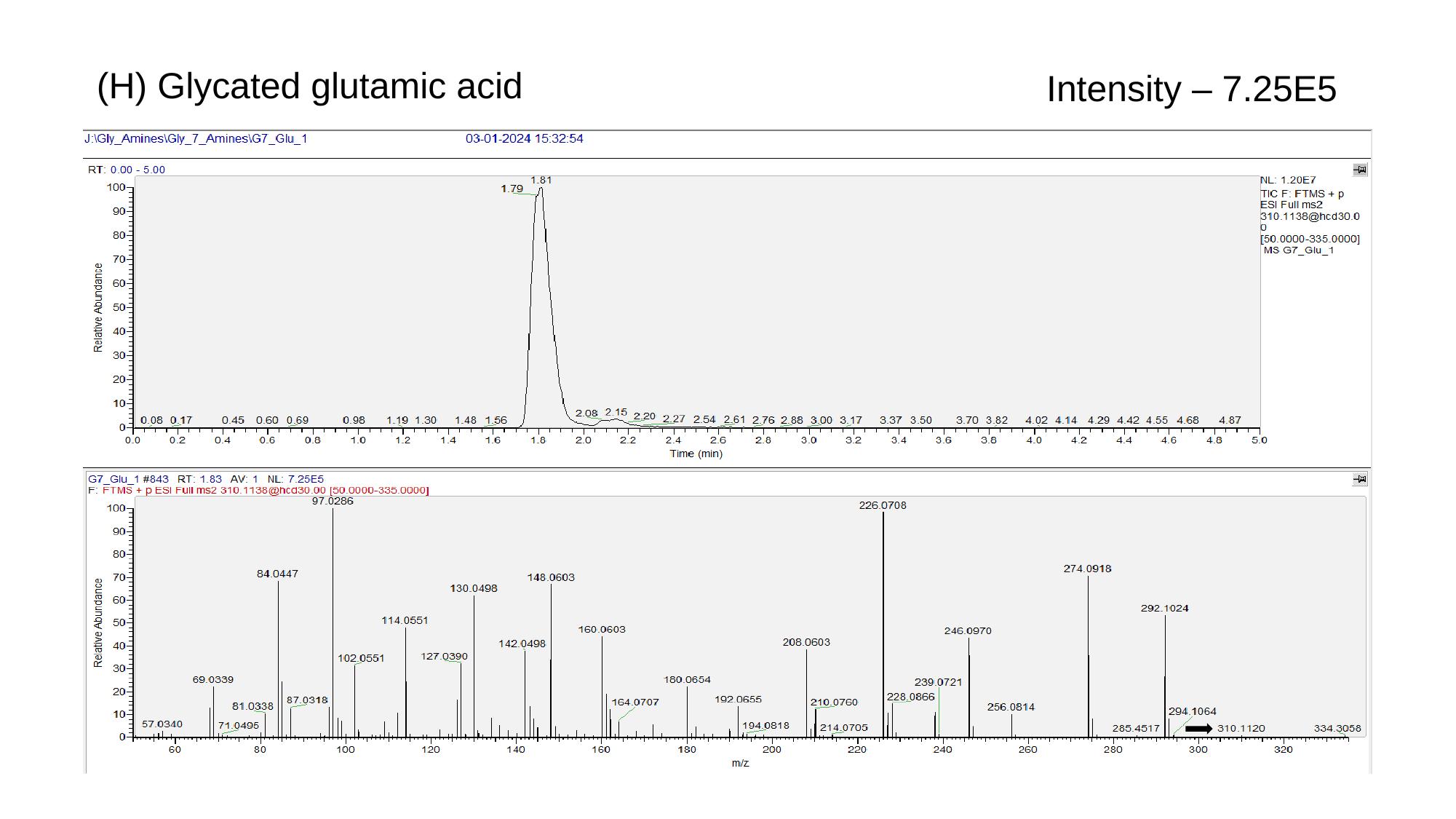

(H) Glycated glutamic acid
Intensity – 7.25E5

### Slide 10
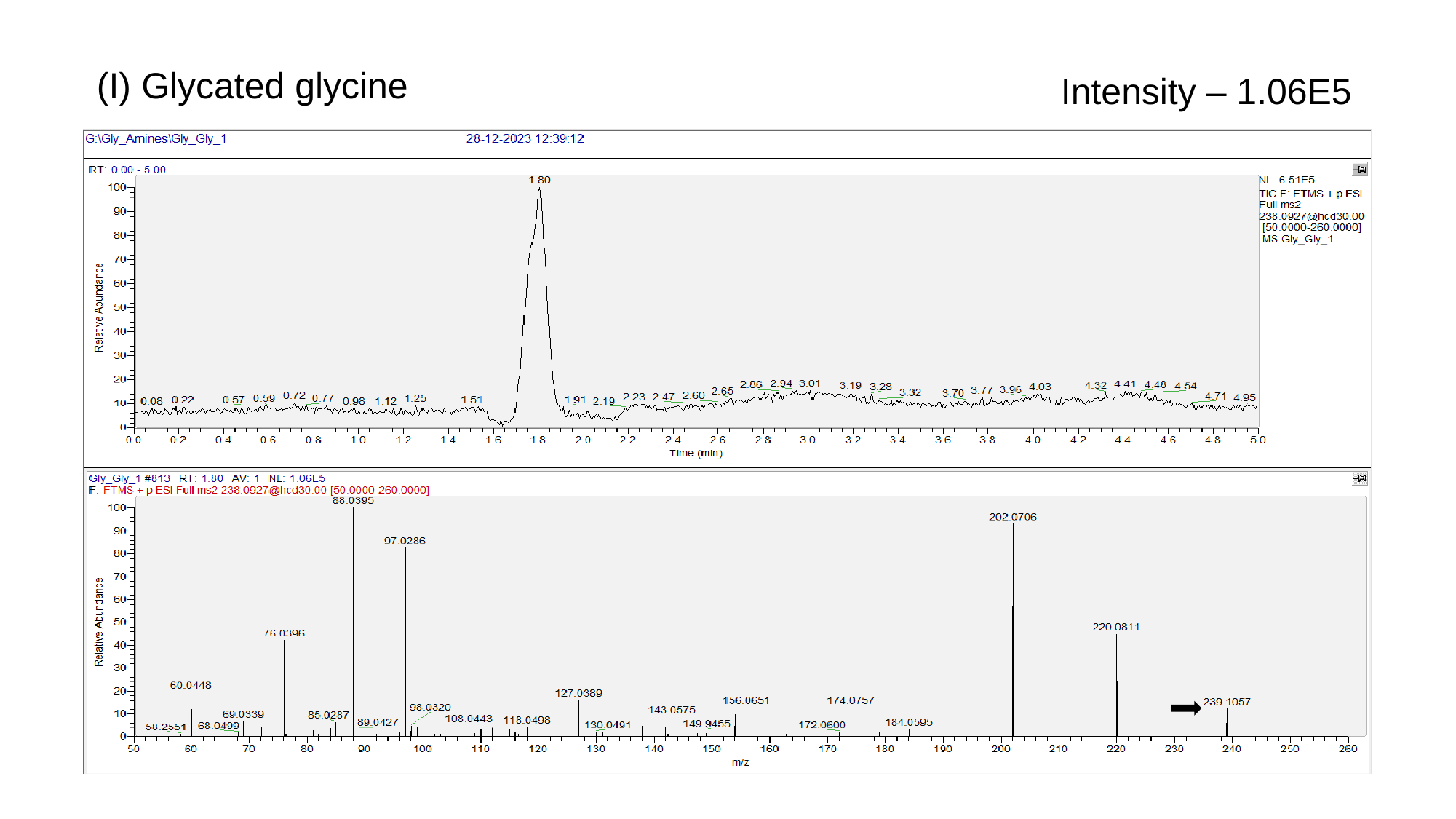

(I) Glycated glycine
Intensity – 1.06E5

### Slide 11
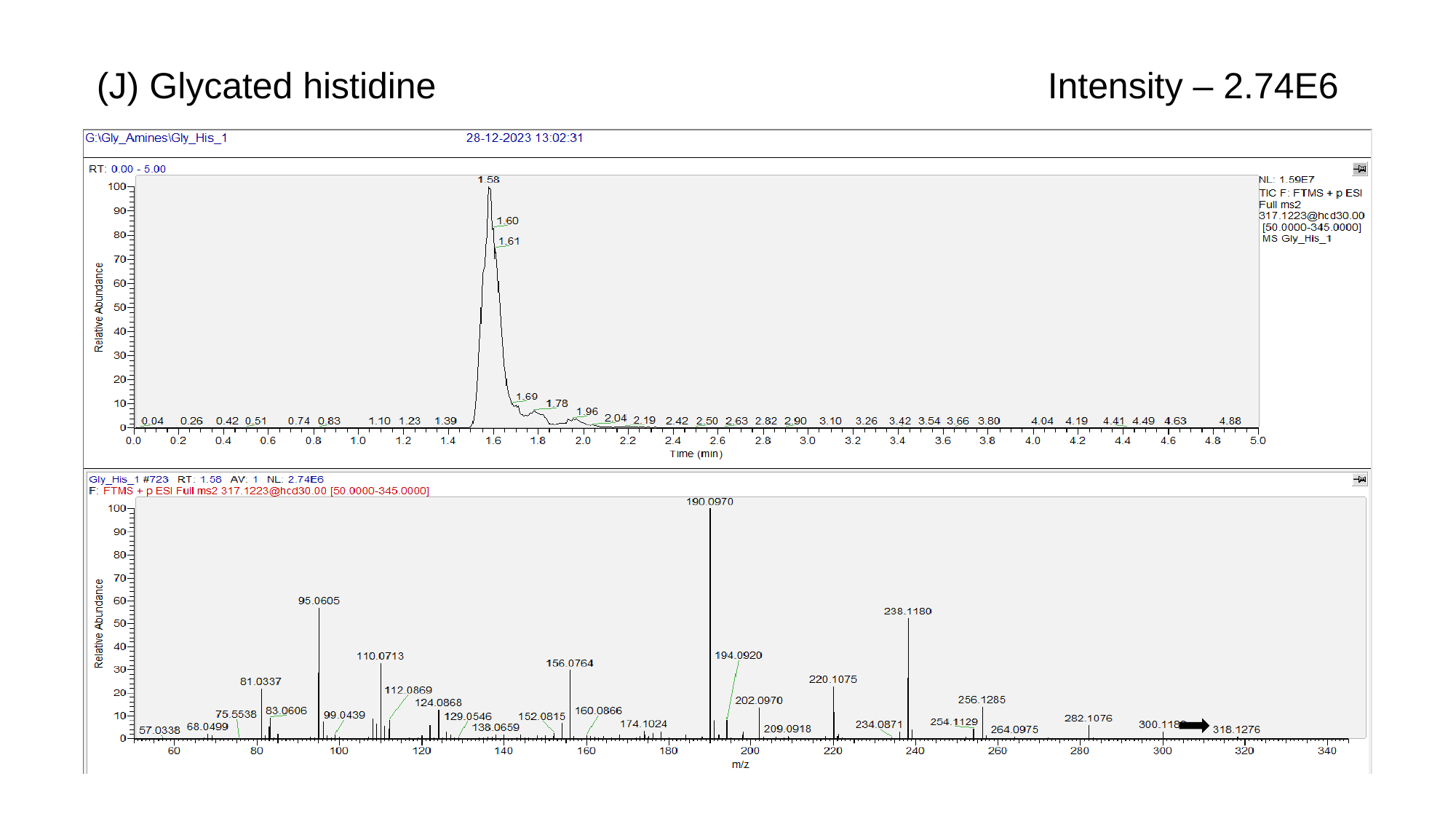

(J) Glycated histidine
Intensity – 2.74E6

### Slide 12
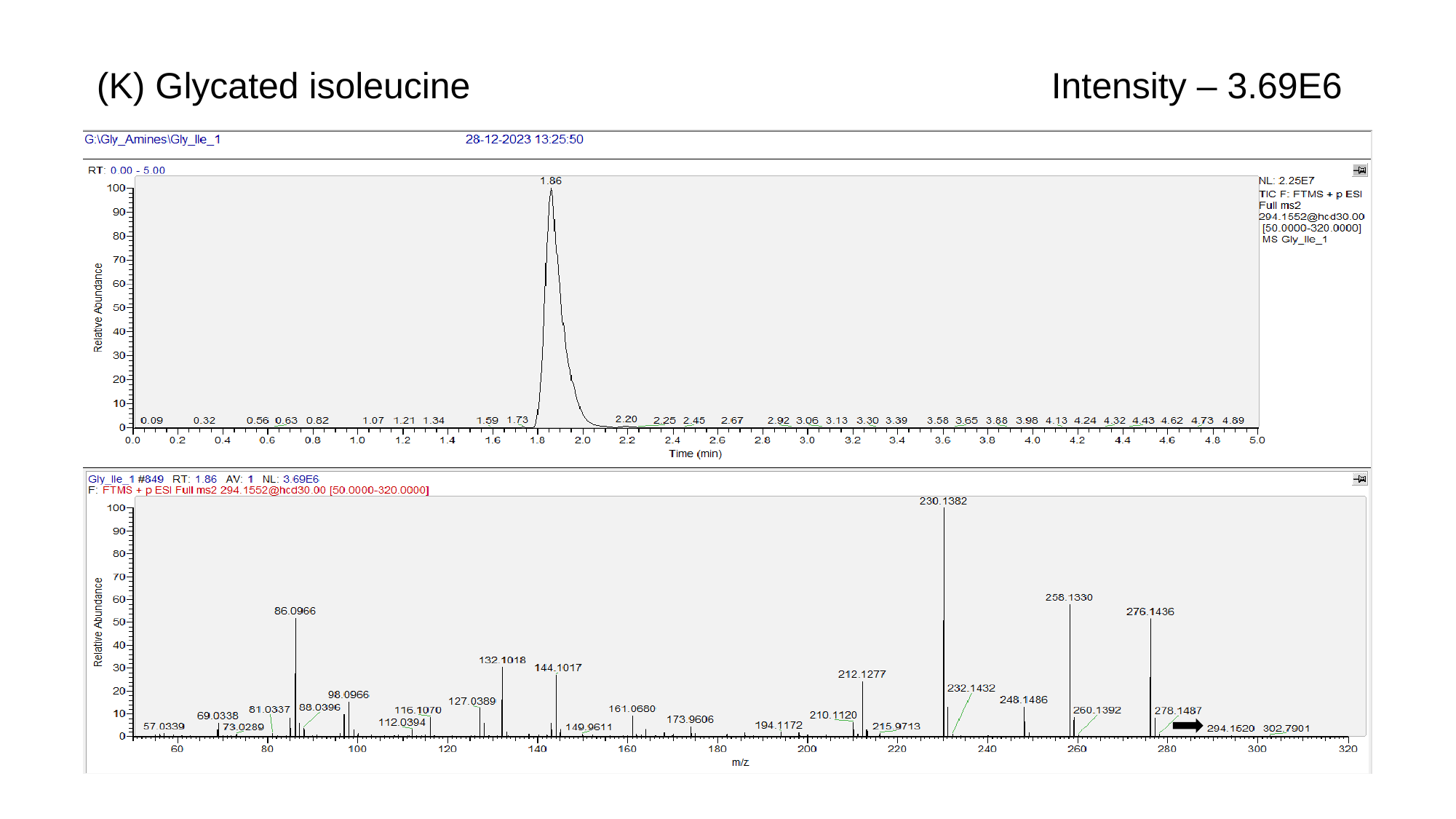

(K) Glycated isoleucine
Intensity – 3.69E6

### Slide 13
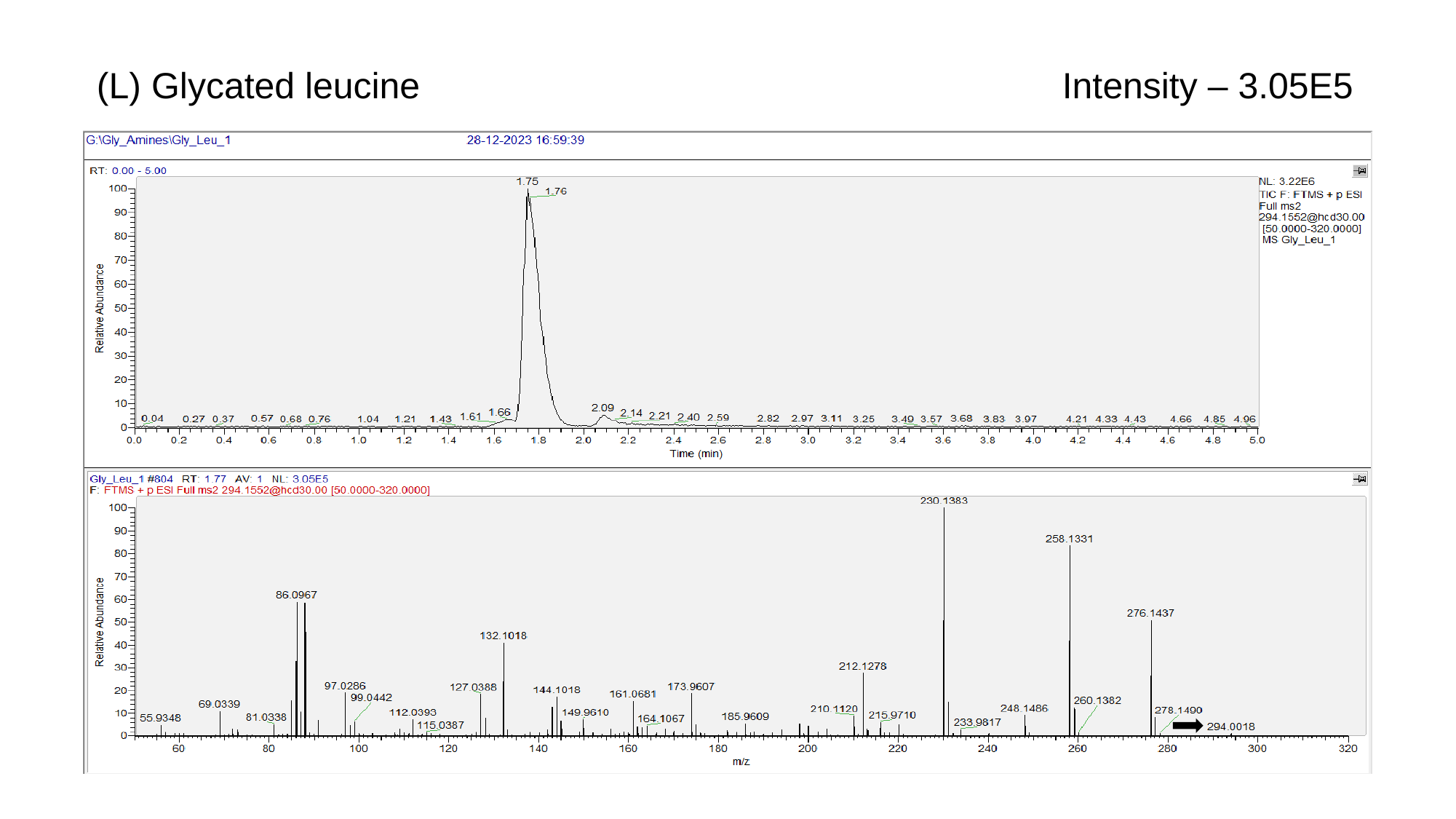

(L) Glycated leucine
Intensity – 3.05E5

### Slide 14
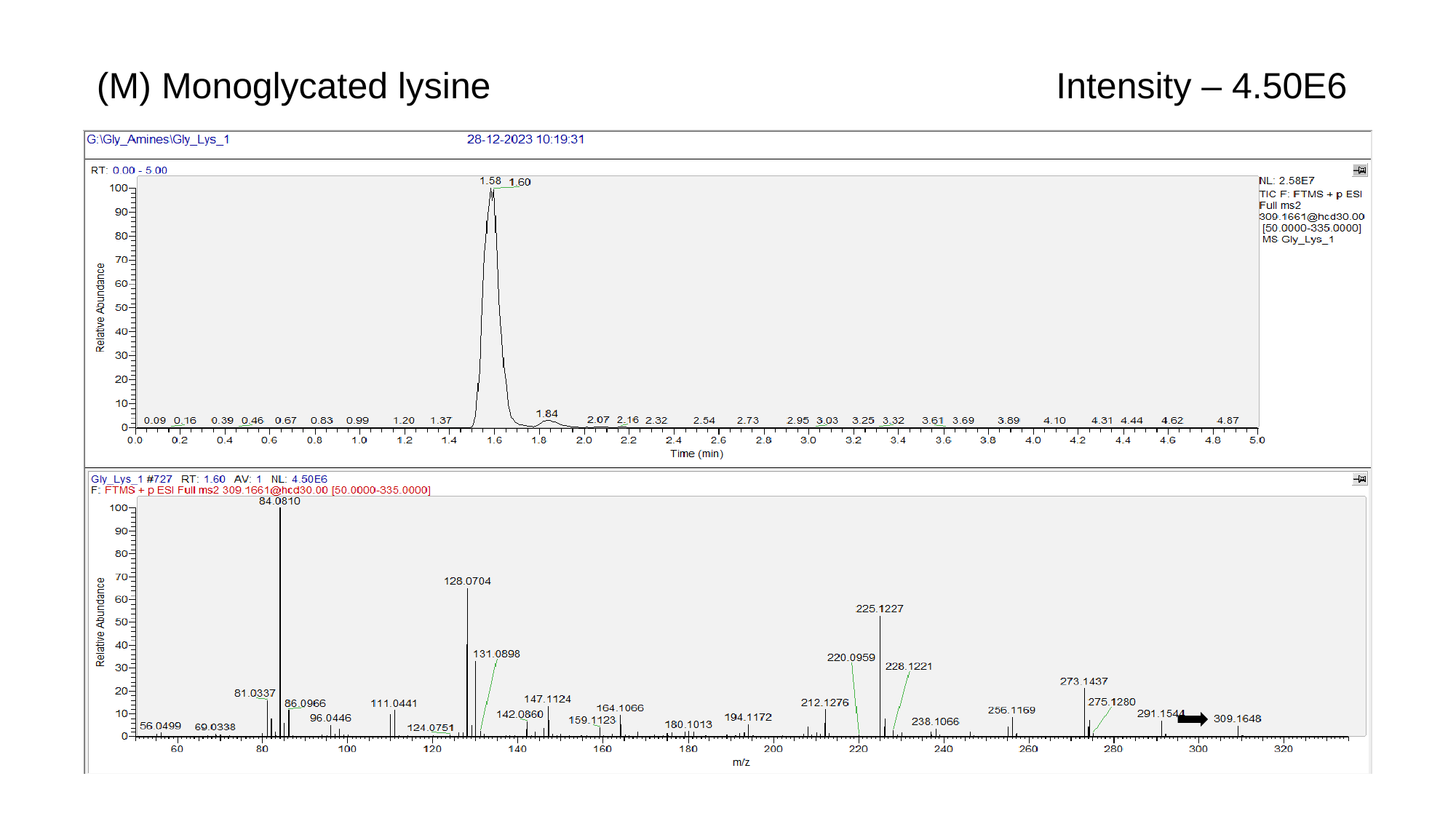

(M) Monoglycated lysine
Intensity – 4.50E6

### Slide 15
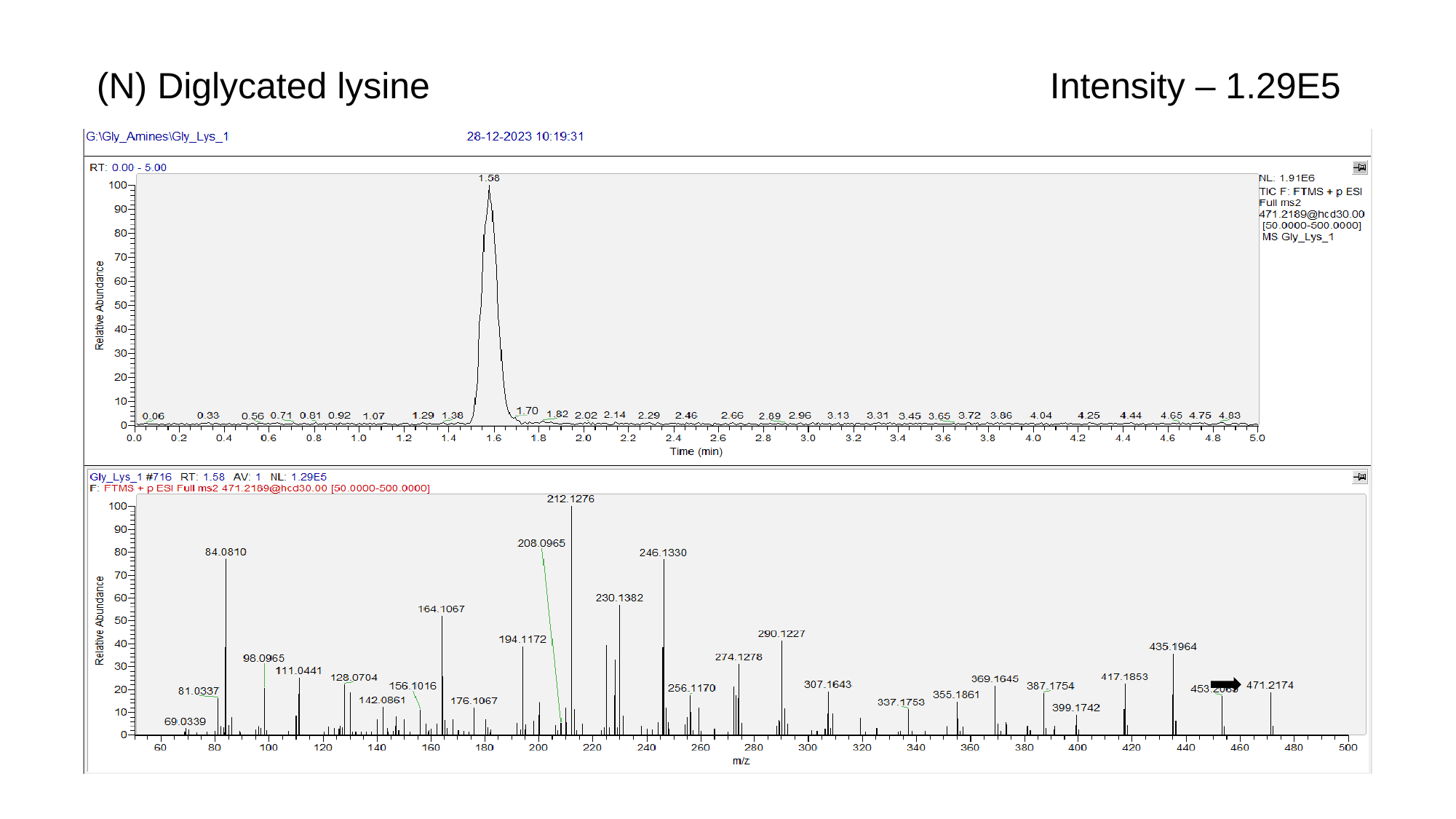

(N) Diglycated lysine
Intensity – 1.29E5

### Slide 16
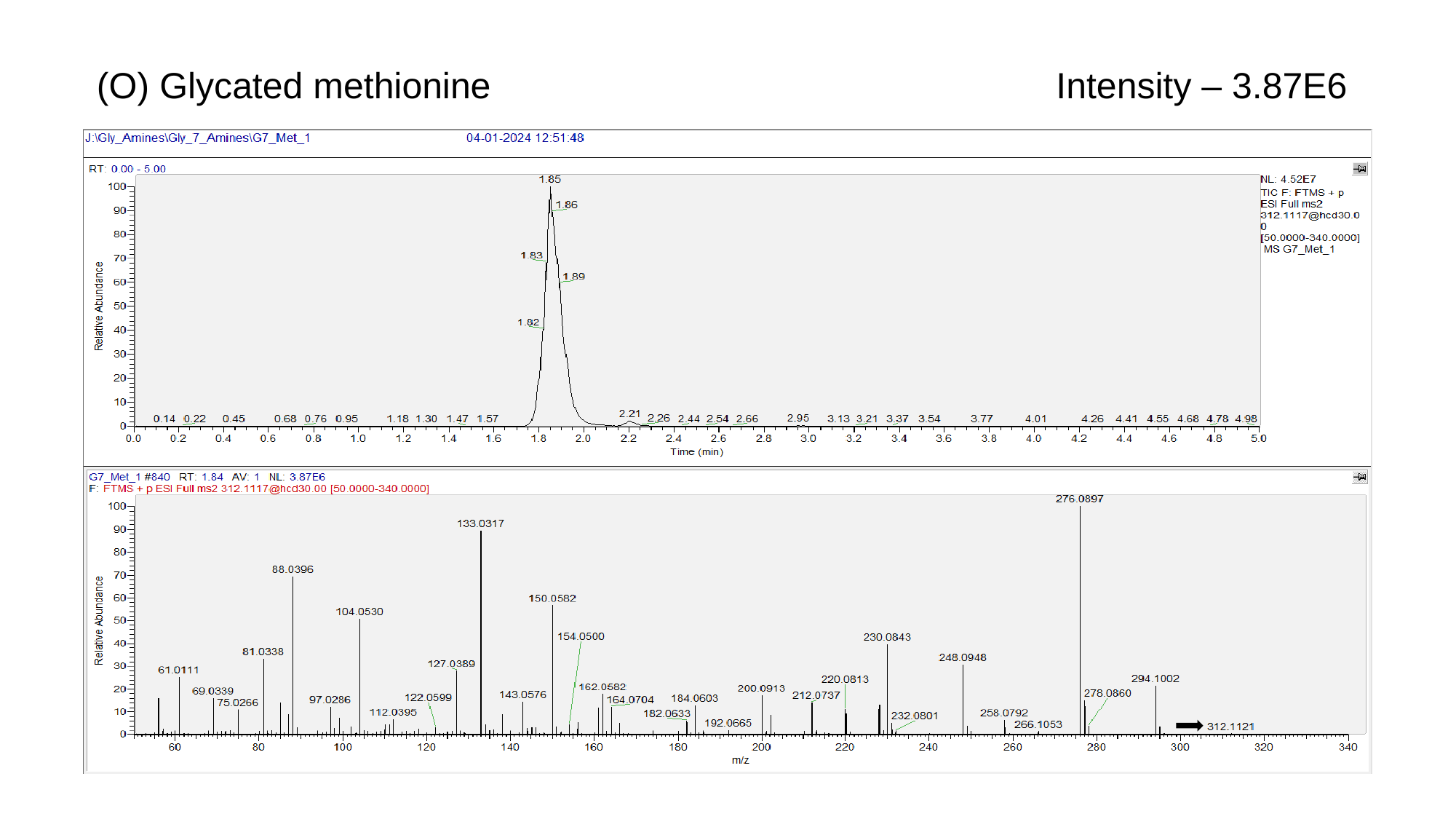

(O) Glycated methionine
Intensity – 3.87E6

### Slide 17
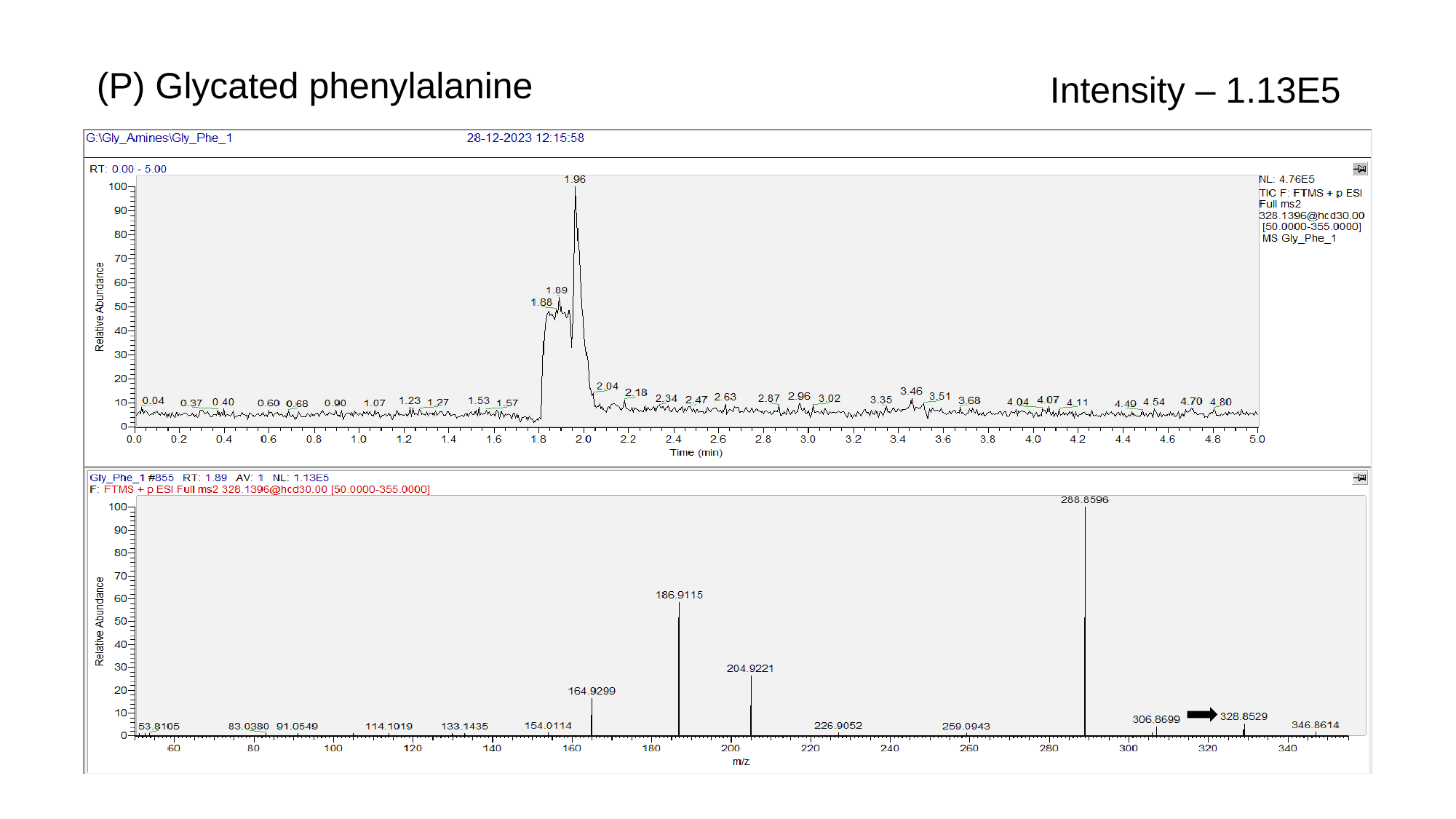

(P) Glycated phenylalanine
Intensity – 1.13E5

### Slide 18
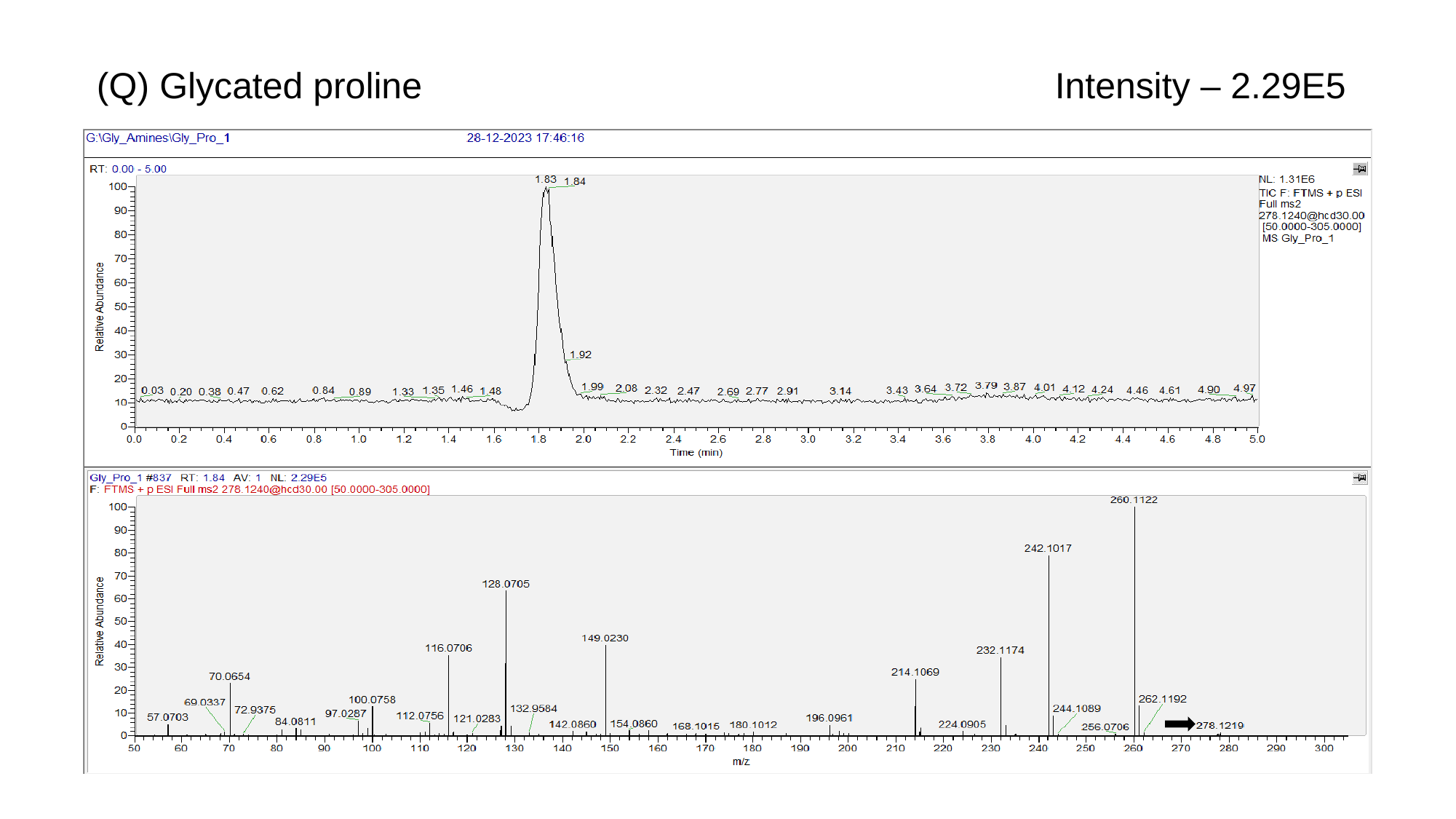

(Q) Glycated proline
Intensity – 2.29E5

### Slide 19
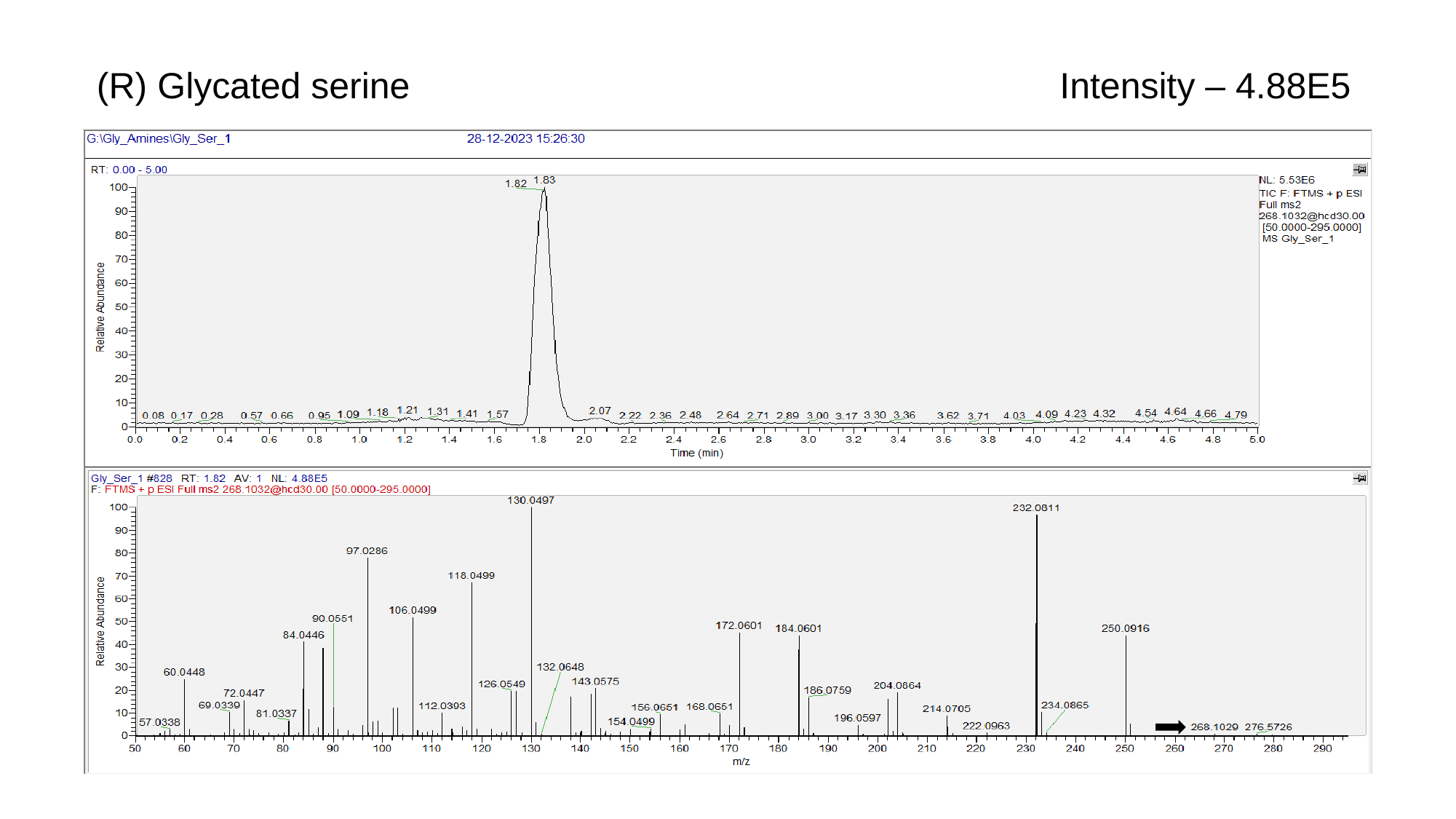

(R) Glycated serine
Intensity – 4.88E5

### Slide 20
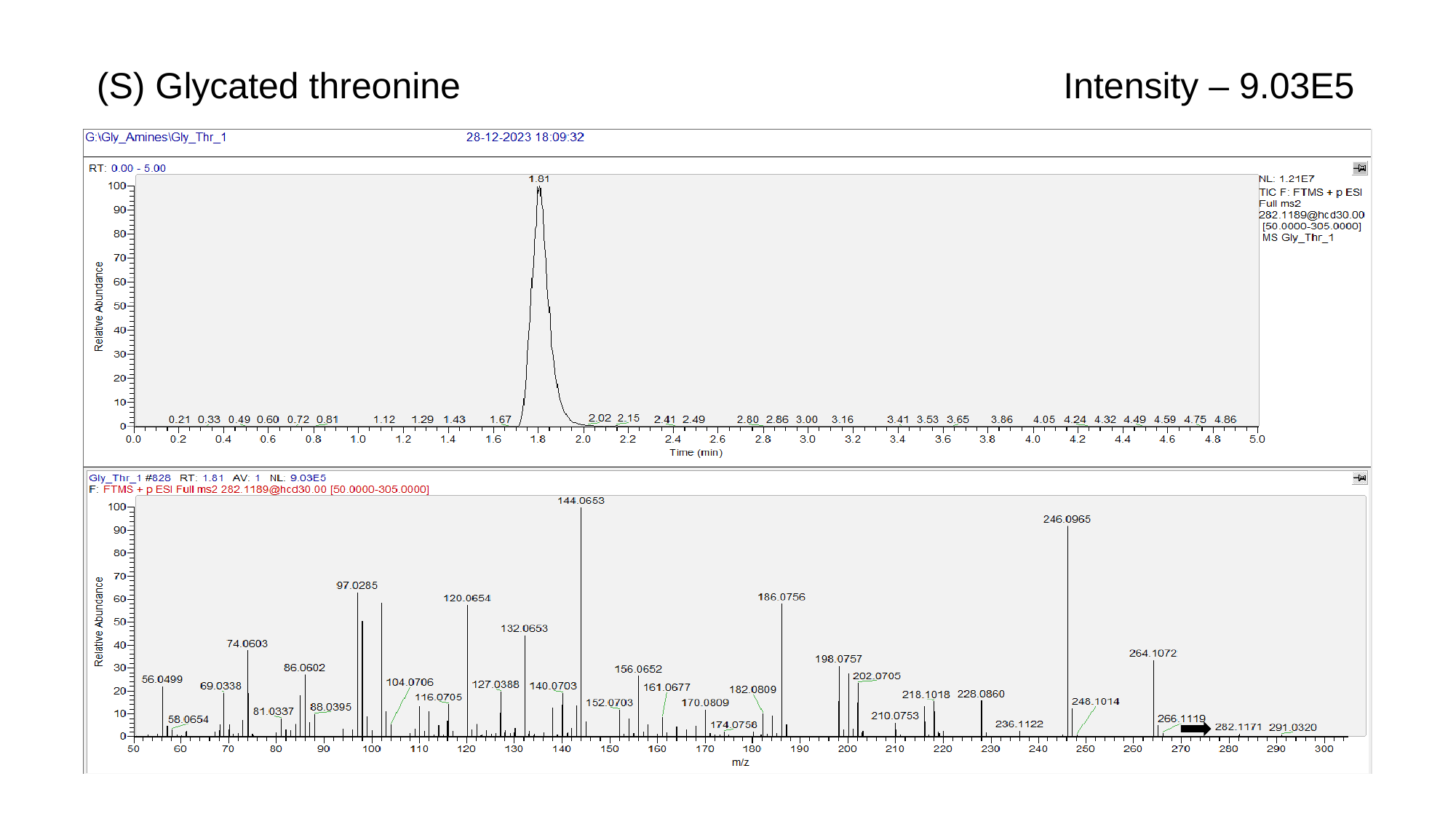

(S) Glycated threonine
Intensity – 9.03E5

### Slide 21
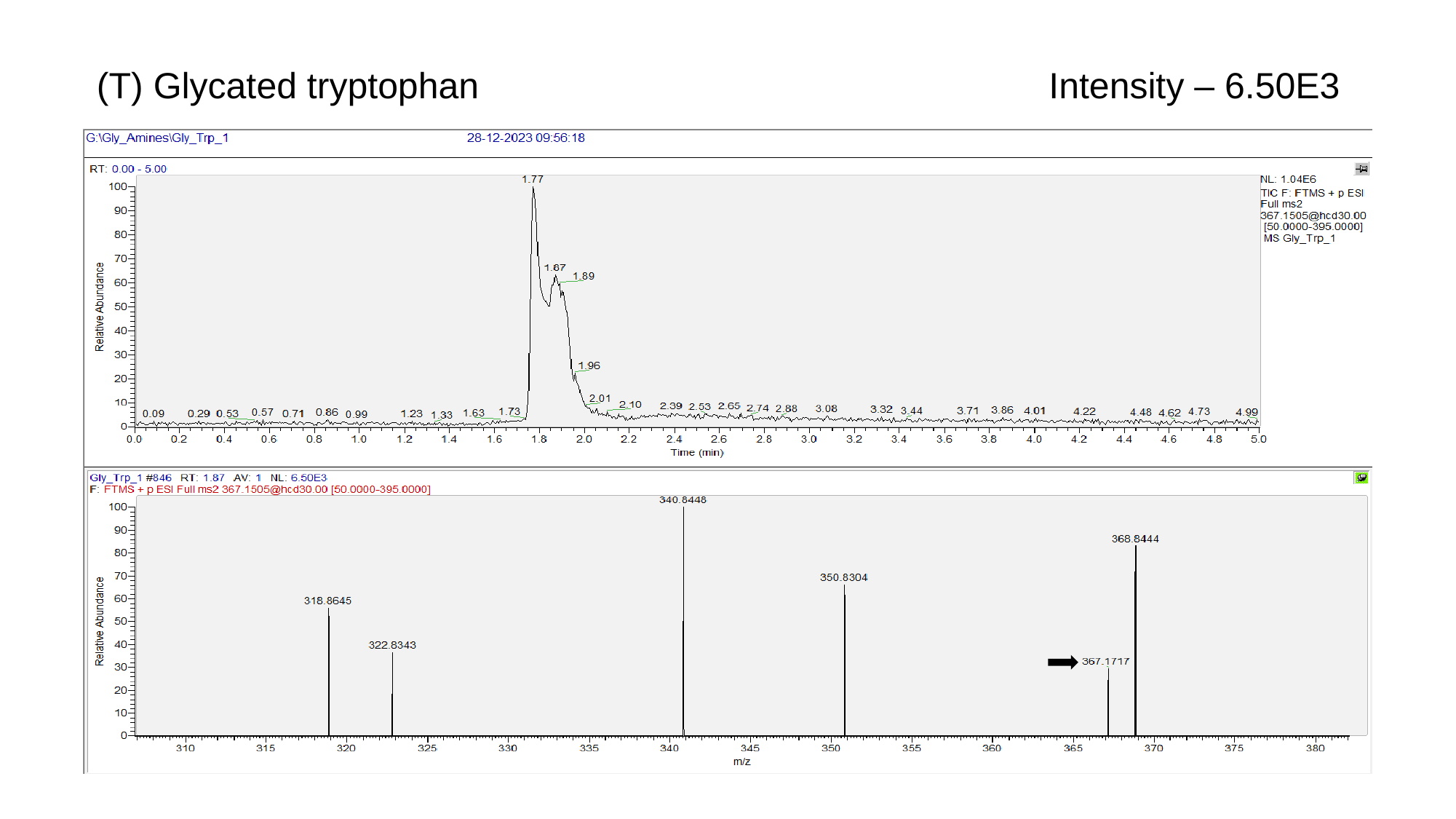

(T) Glycated tryptophan
Intensity – 6.50E3

### Slide 22
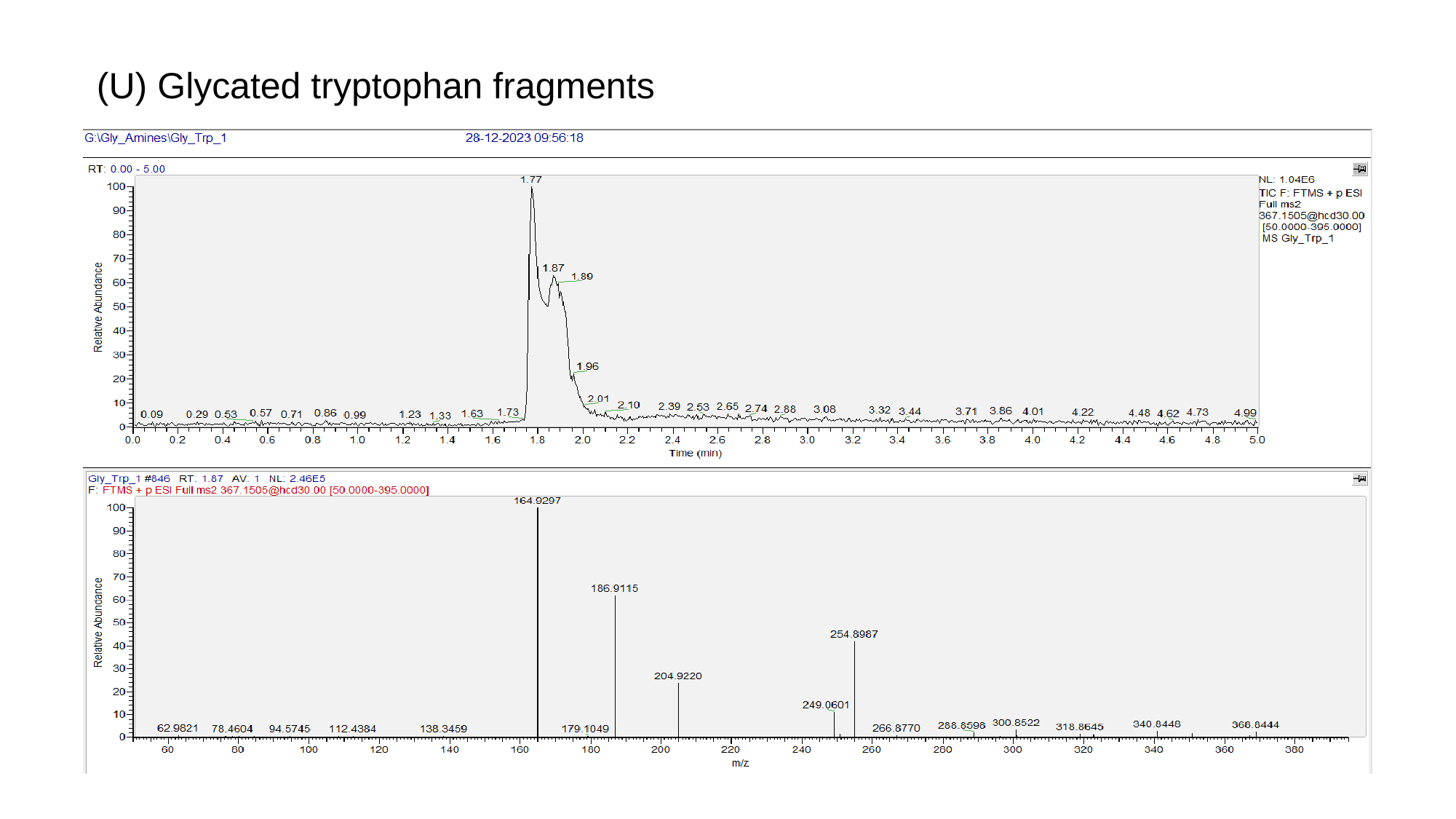

(U) Glycated tryptophan fragments

### Slide 23
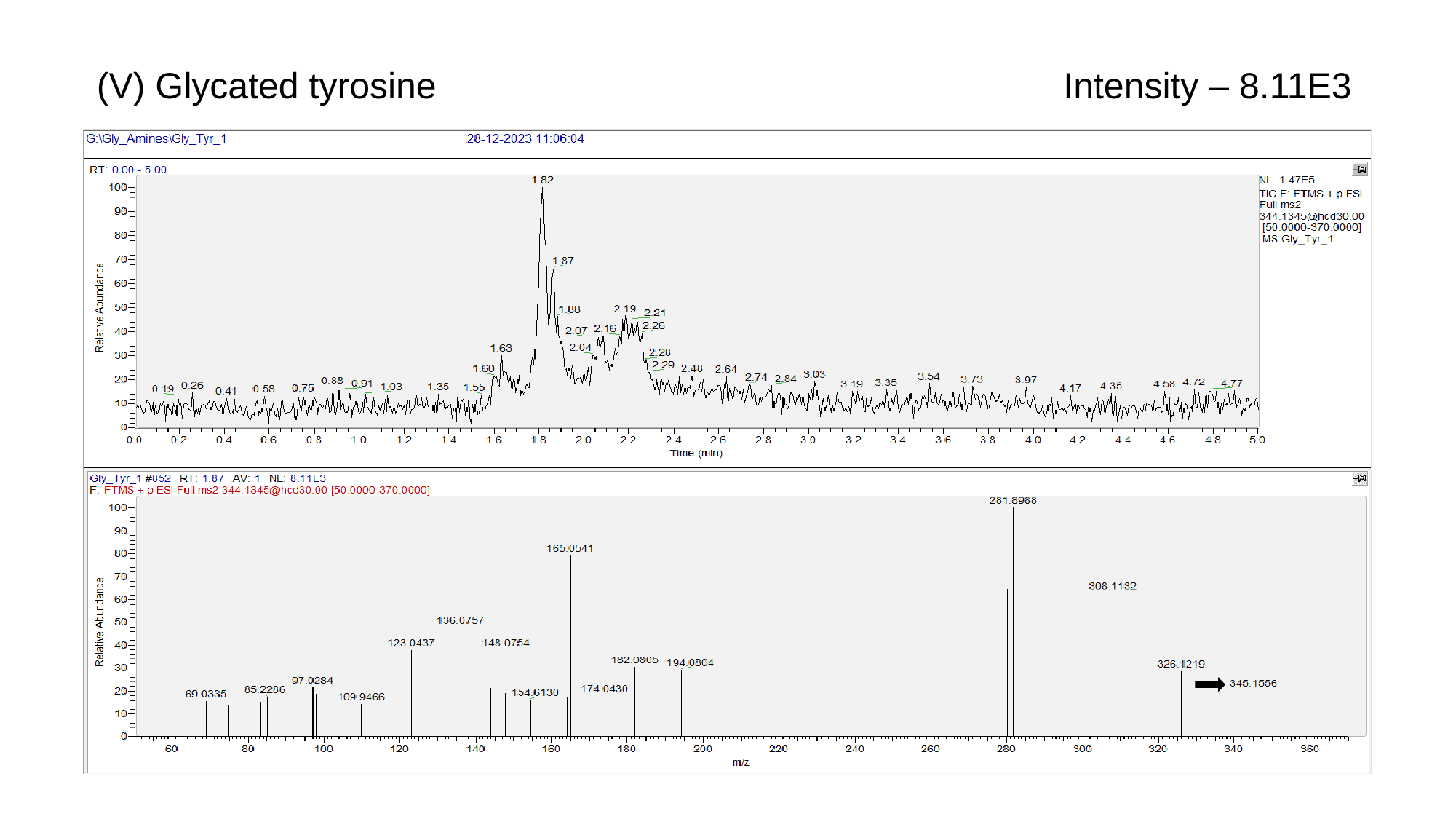

(V) Glycated tyrosine
Intensity – 8.11E3

### Slide 24
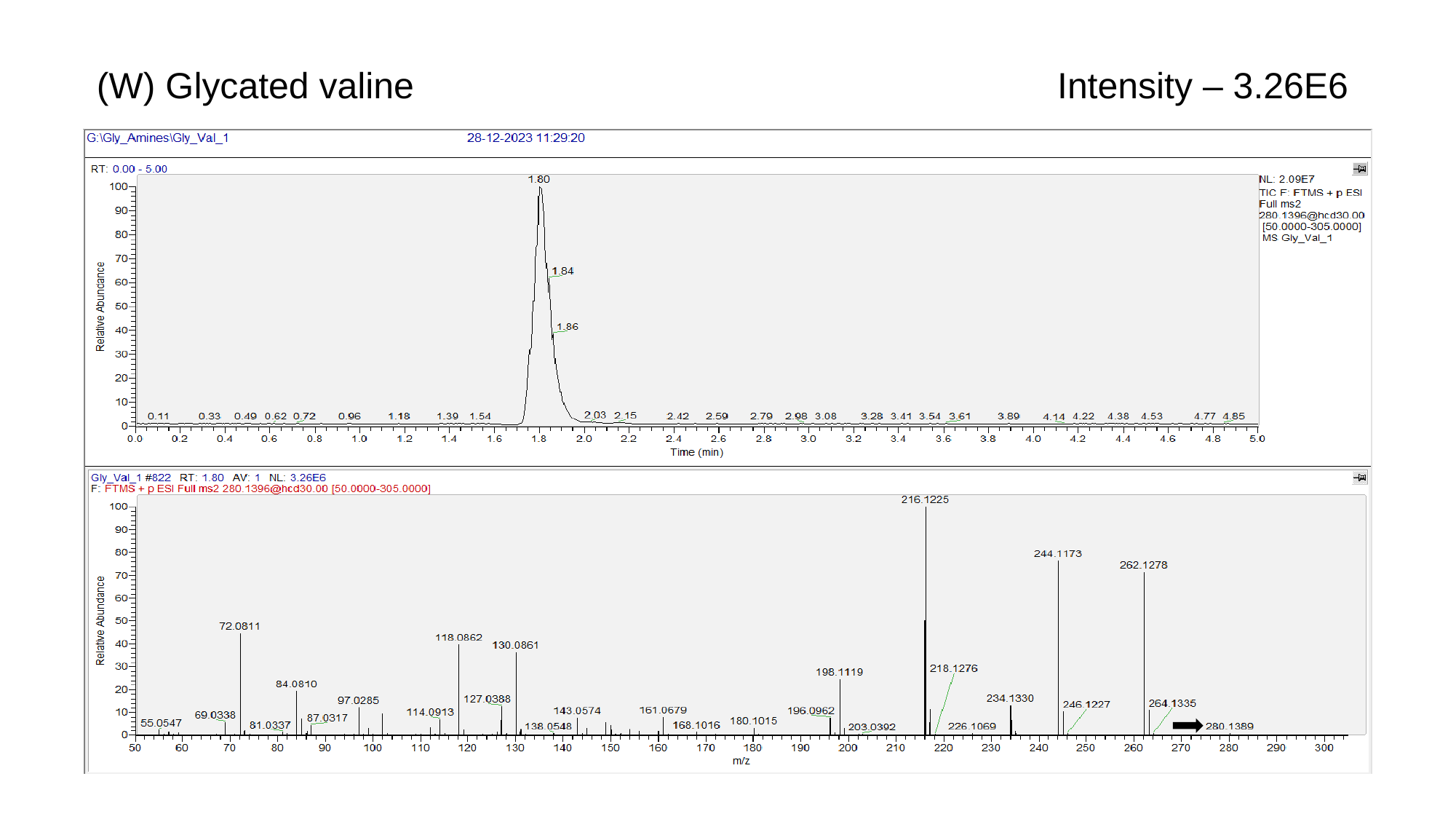

(W) Glycated valine
Intensity – 3.26E6

### Slide 25
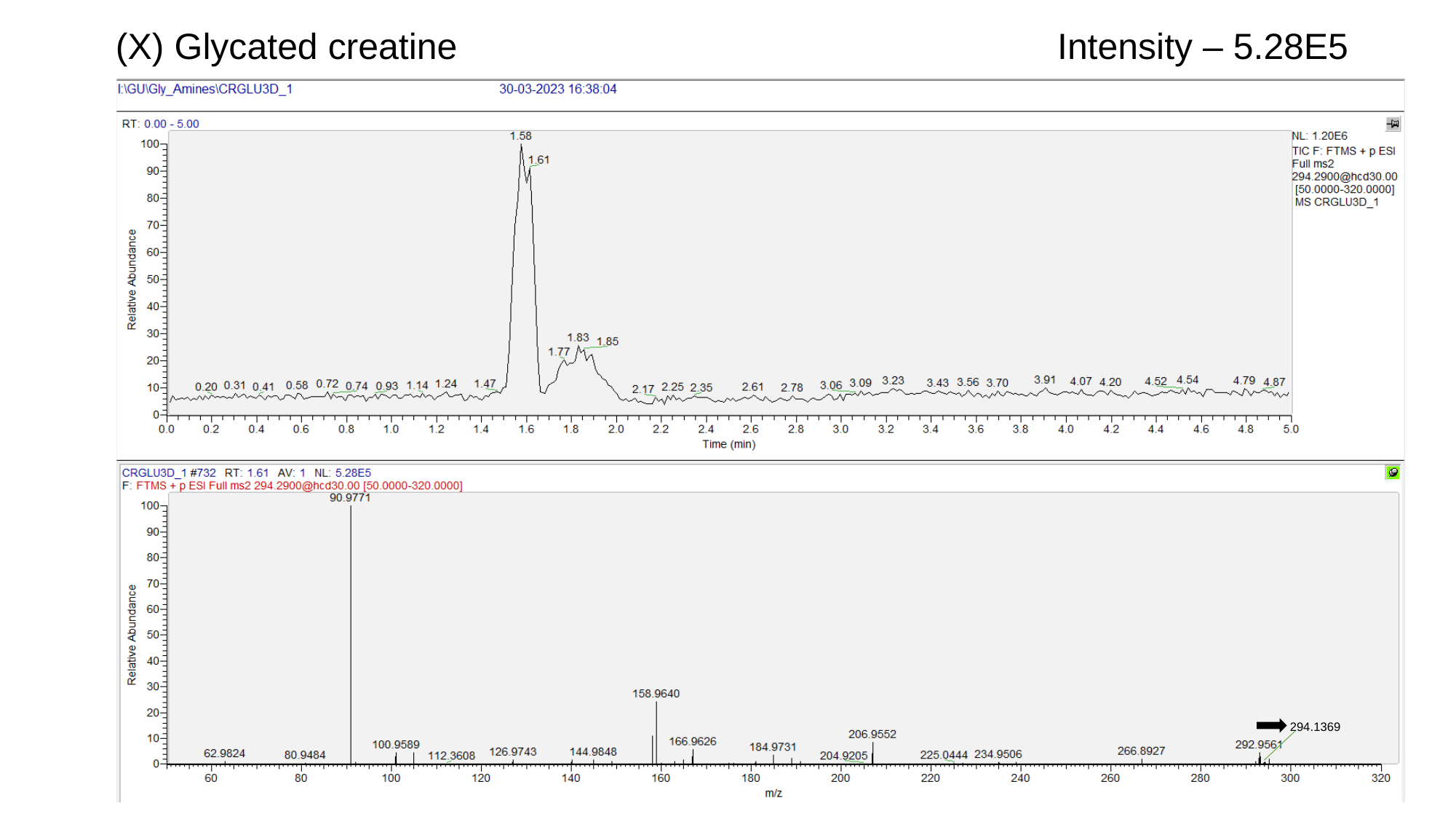

(X) Glycated creatine
Intensity – 5.28E5
294.1369

### Slide 26
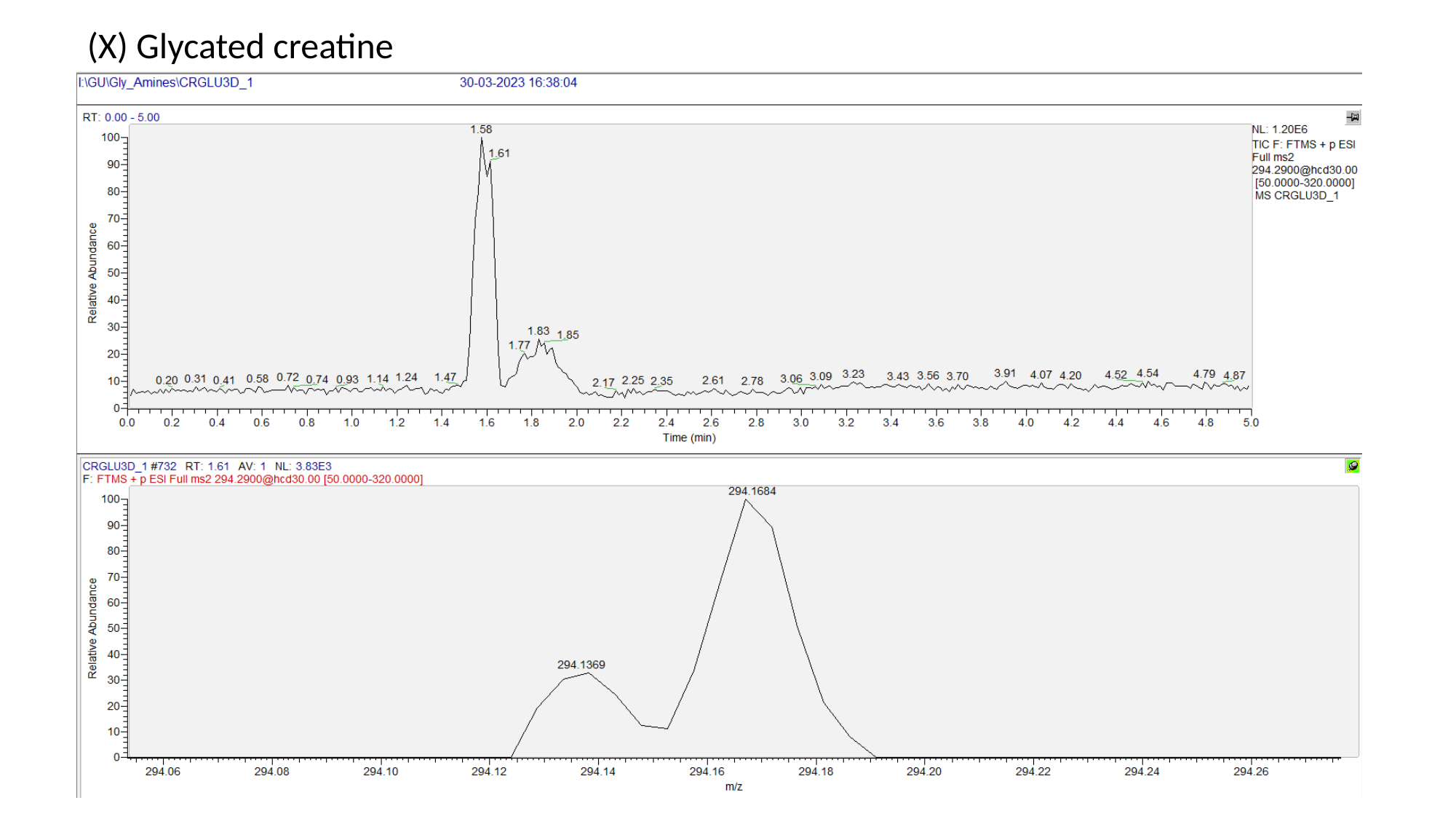

(X) Glycated creatine

### Slide 27
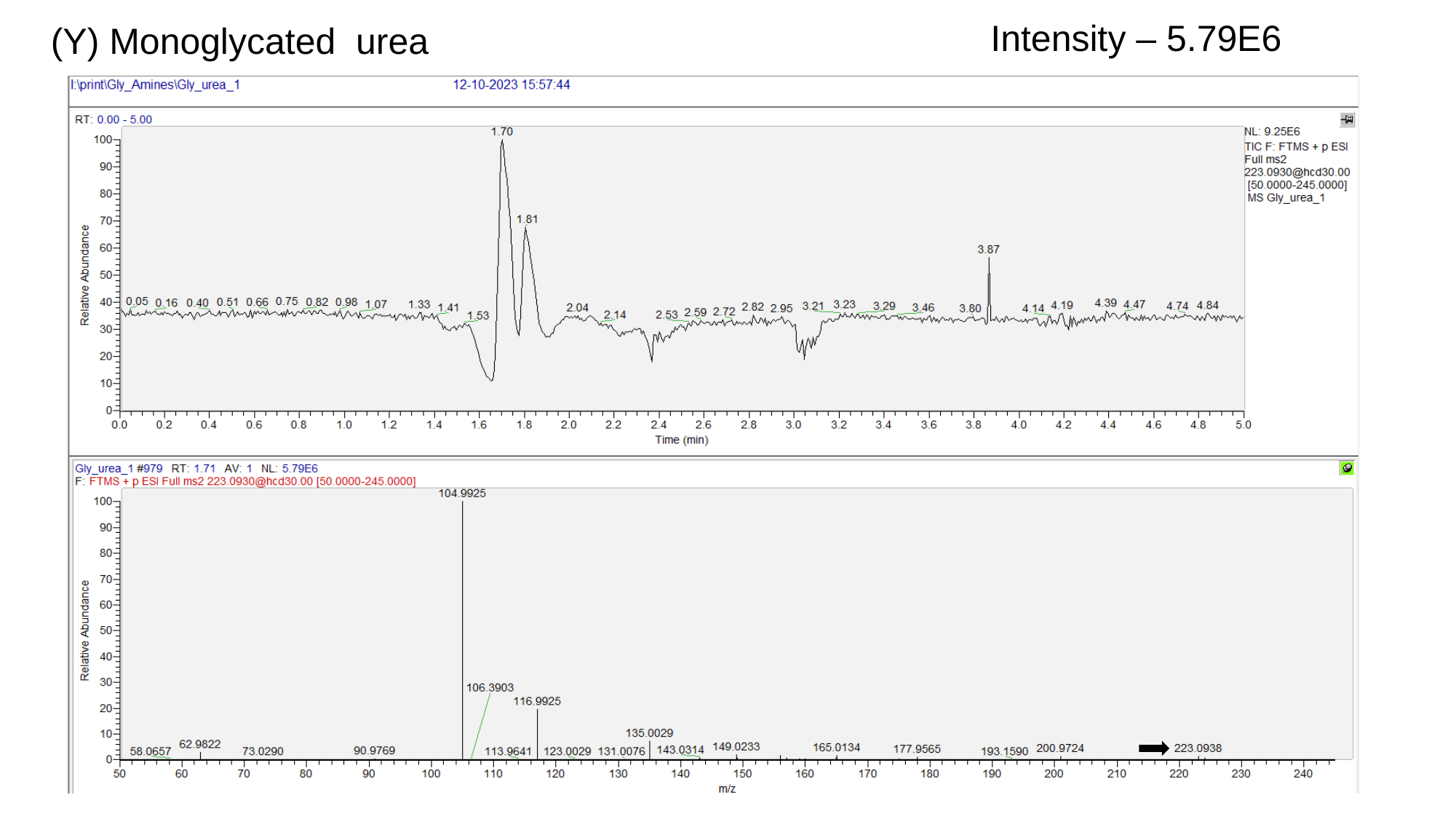

(Y) Monoglycated urea
Intensity – 5.79E6

### Slide 28
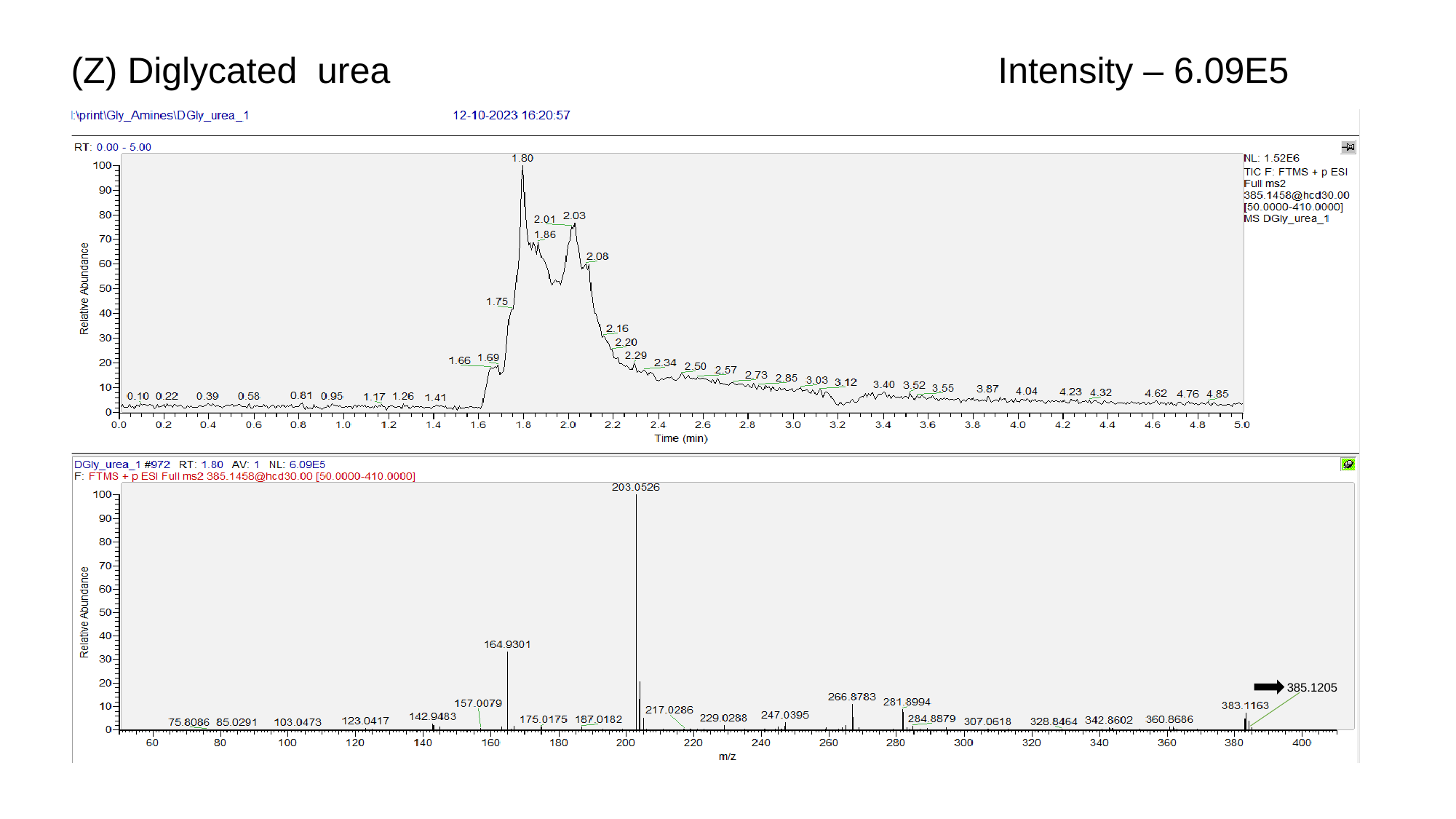

Intensity – 6.09E5
(Z) Diglycated urea
385.1205

### Slide 29
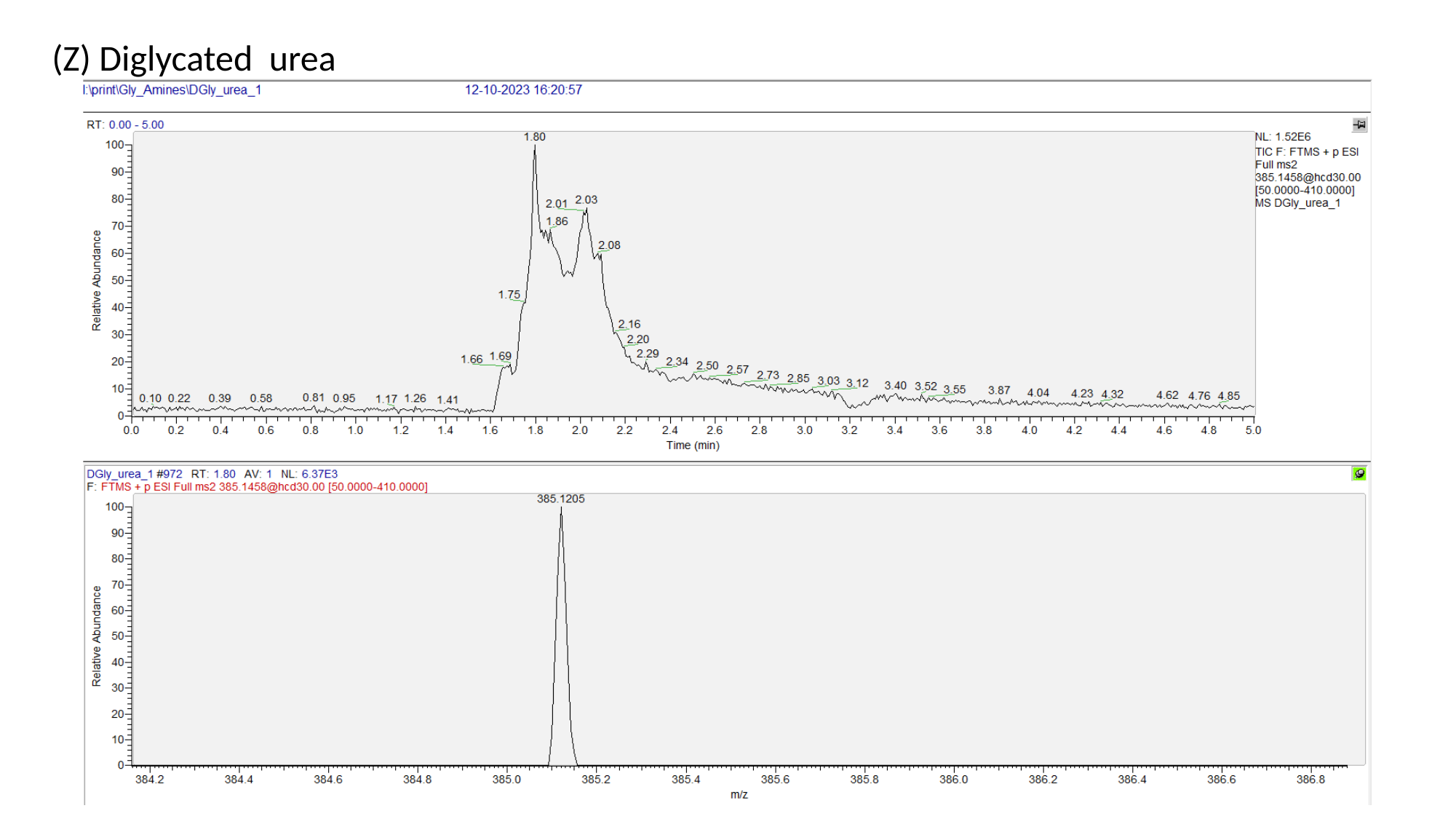

(Z) Diglycated urea

### Slide 30
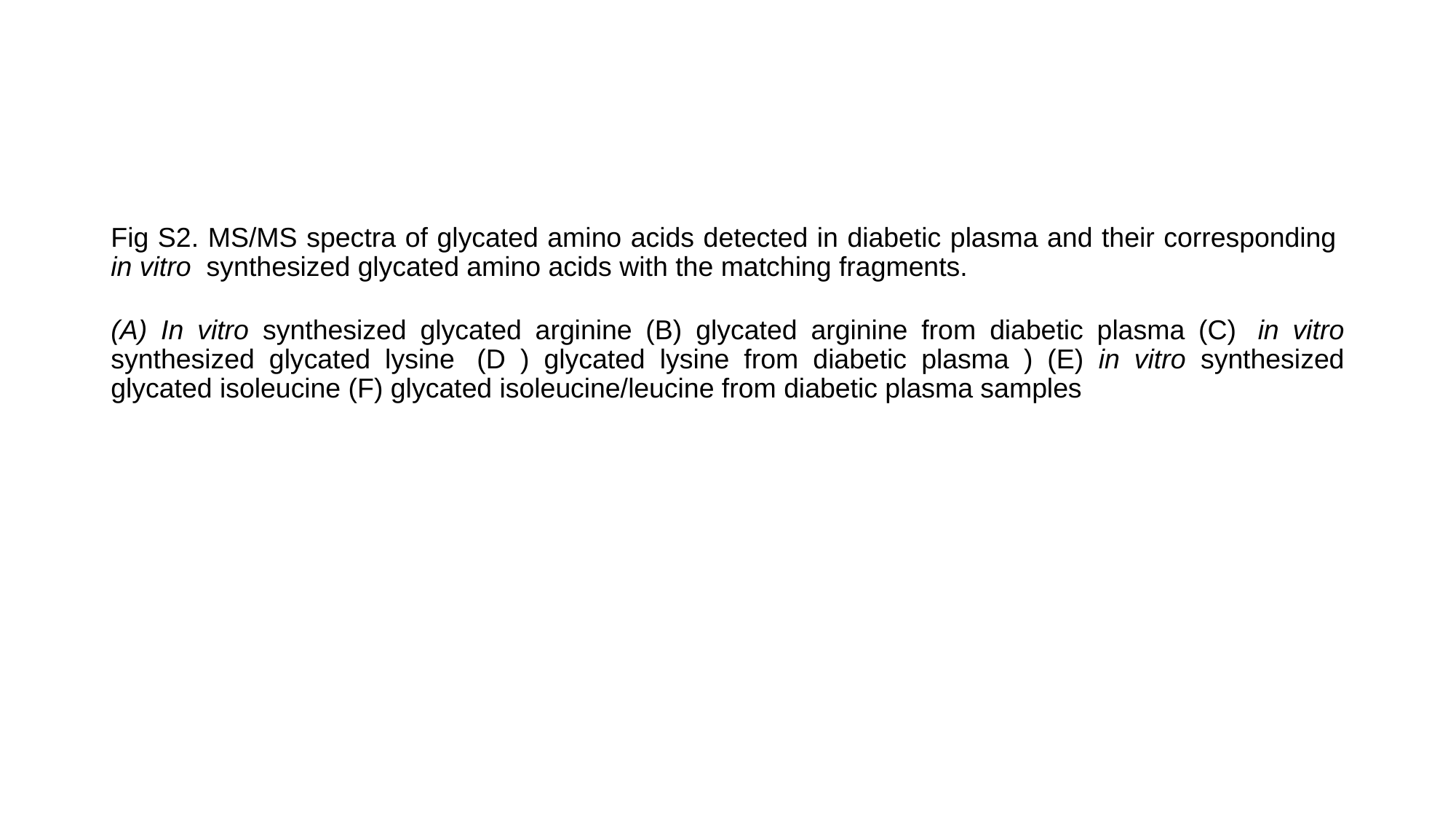

Fig S2. MS/MS spectra of glycated amino acids detected in diabetic plasma and their corresponding  in vitro  synthesized glycated amino acids with the matching fragments.
(A) In vitro synthesized glycated arginine (B) glycated arginine from diabetic plasma (C)  in vitro synthesized glycated lysine  (D ) glycated lysine from diabetic plasma ) (E) in vitro synthesized glycated isoleucine (F) glycated isoleucine/leucine from diabetic plasma samples

### Slide 31
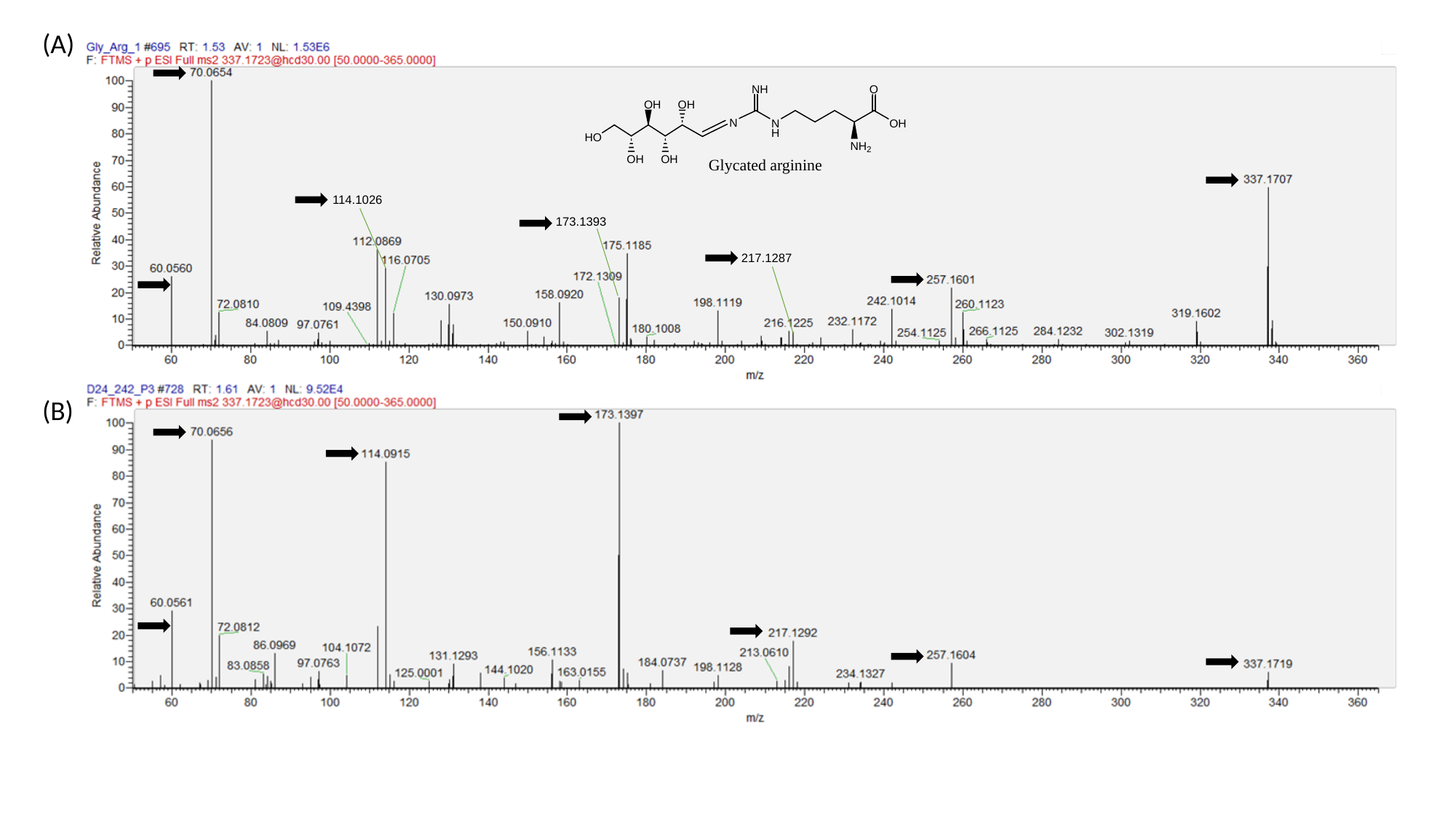

(A)
114.1026
173.1393
217.1287
(B)

### Slide 32
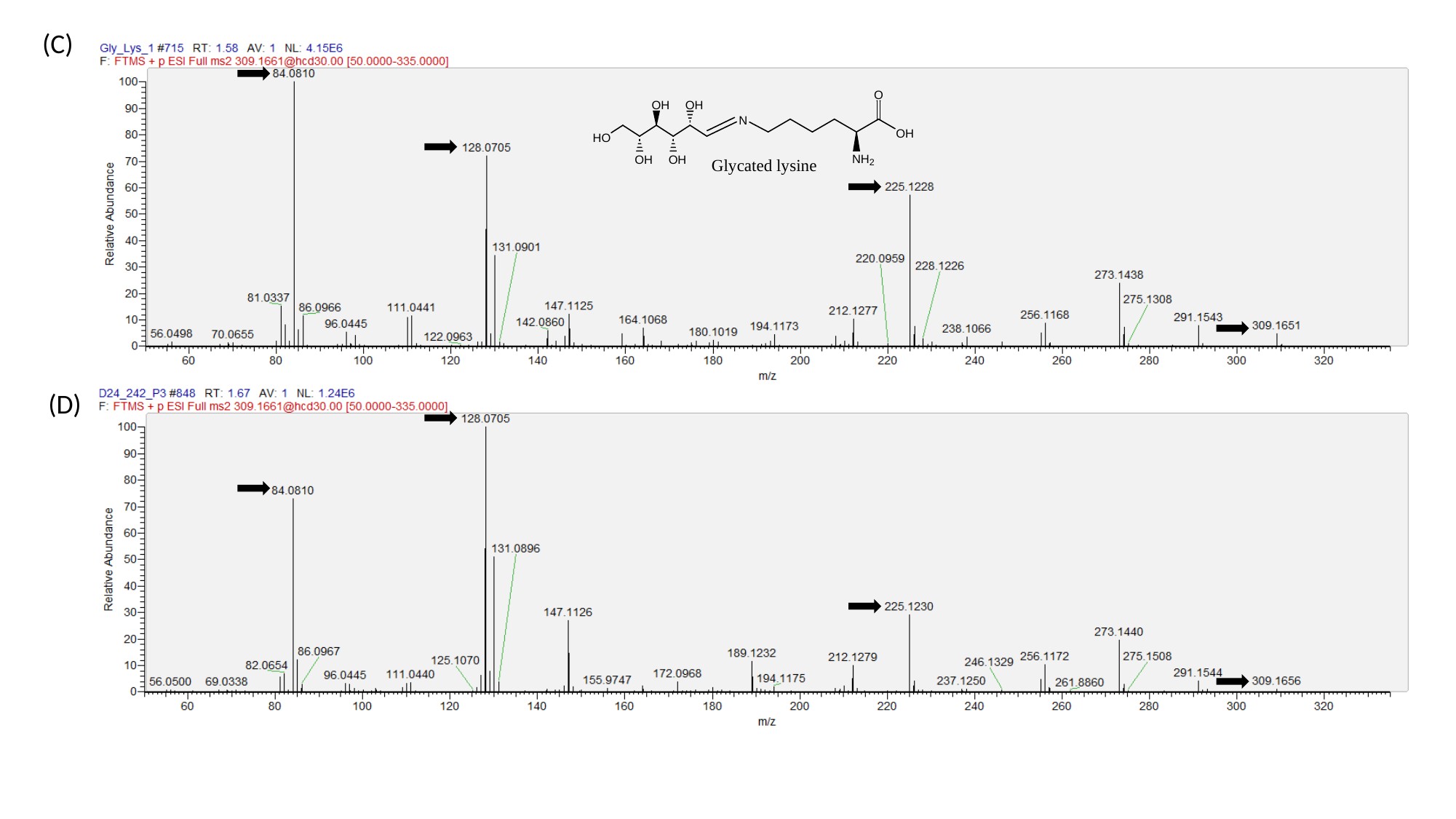

(C)
(D)

### Slide 33
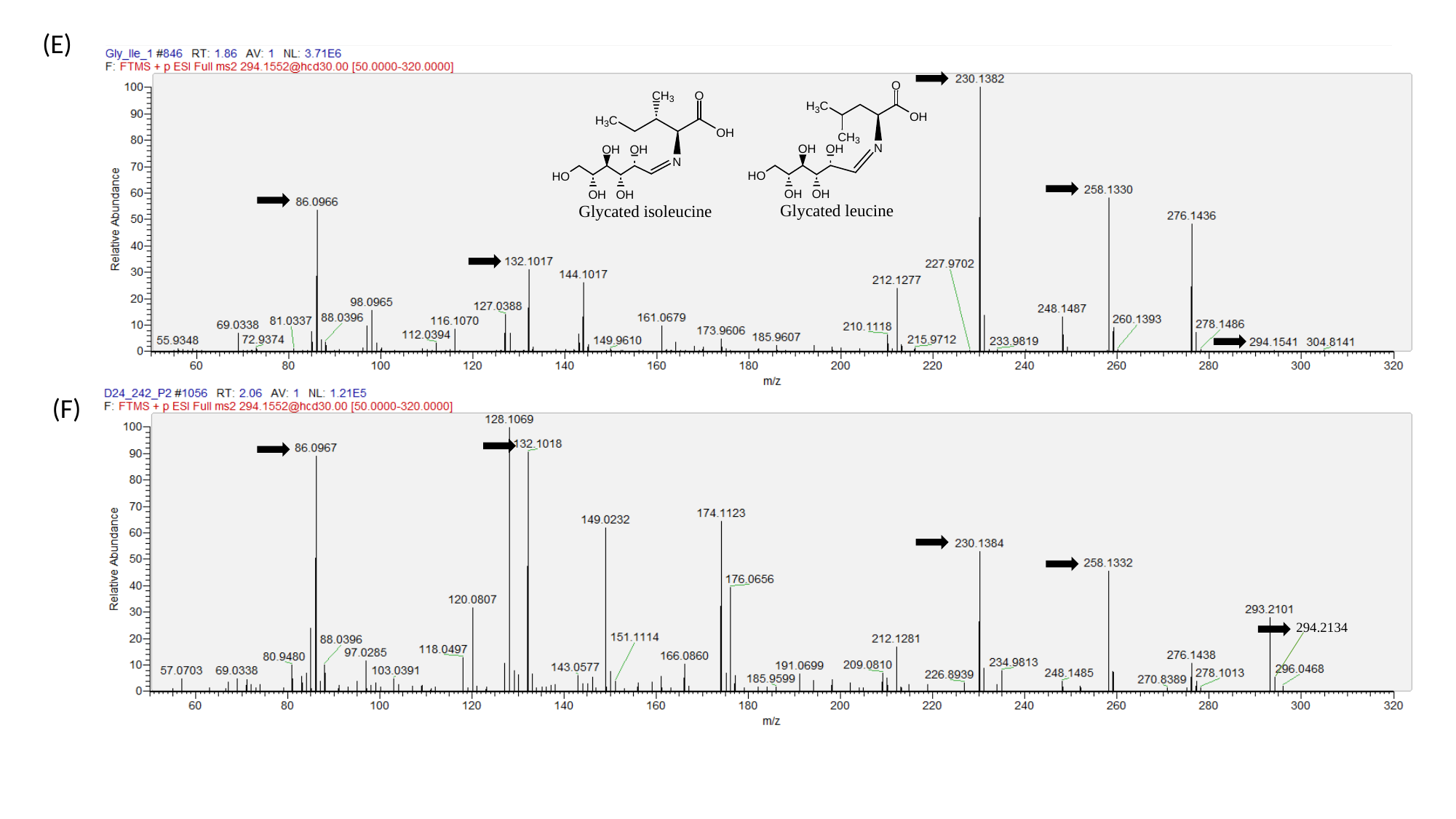

(E)
294.2134
(F)

### Slide 34
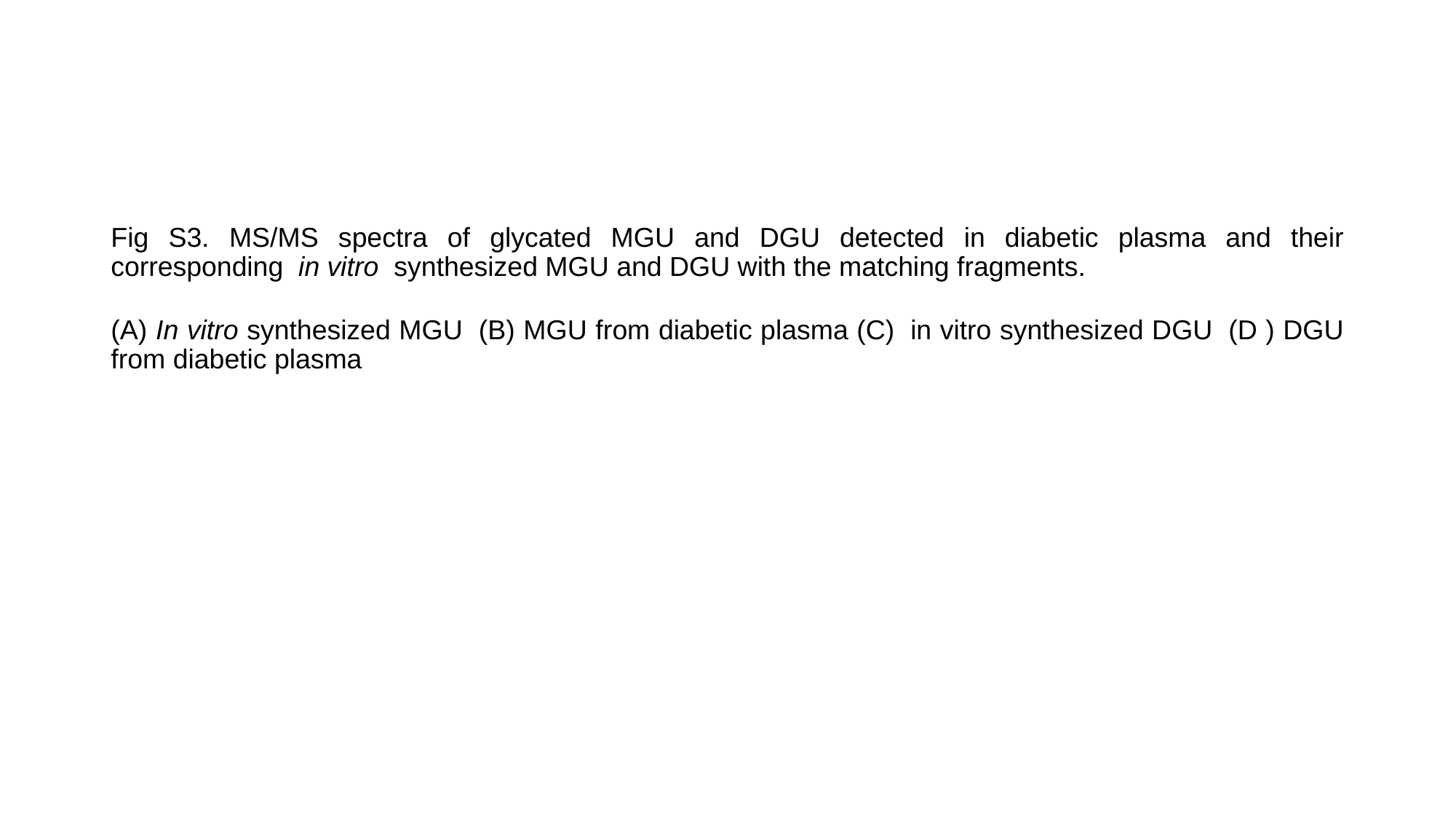

Fig S3. MS/MS spectra of glycated MGU and DGU detected in diabetic plasma and their corresponding  in vitro  synthesized MGU and DGU with the matching fragments.
(A) In vitro synthesized MGU  (B) MGU from diabetic plasma (C)  in vitro synthesized DGU  (D ) DGU from diabetic plasma

### Slide 35
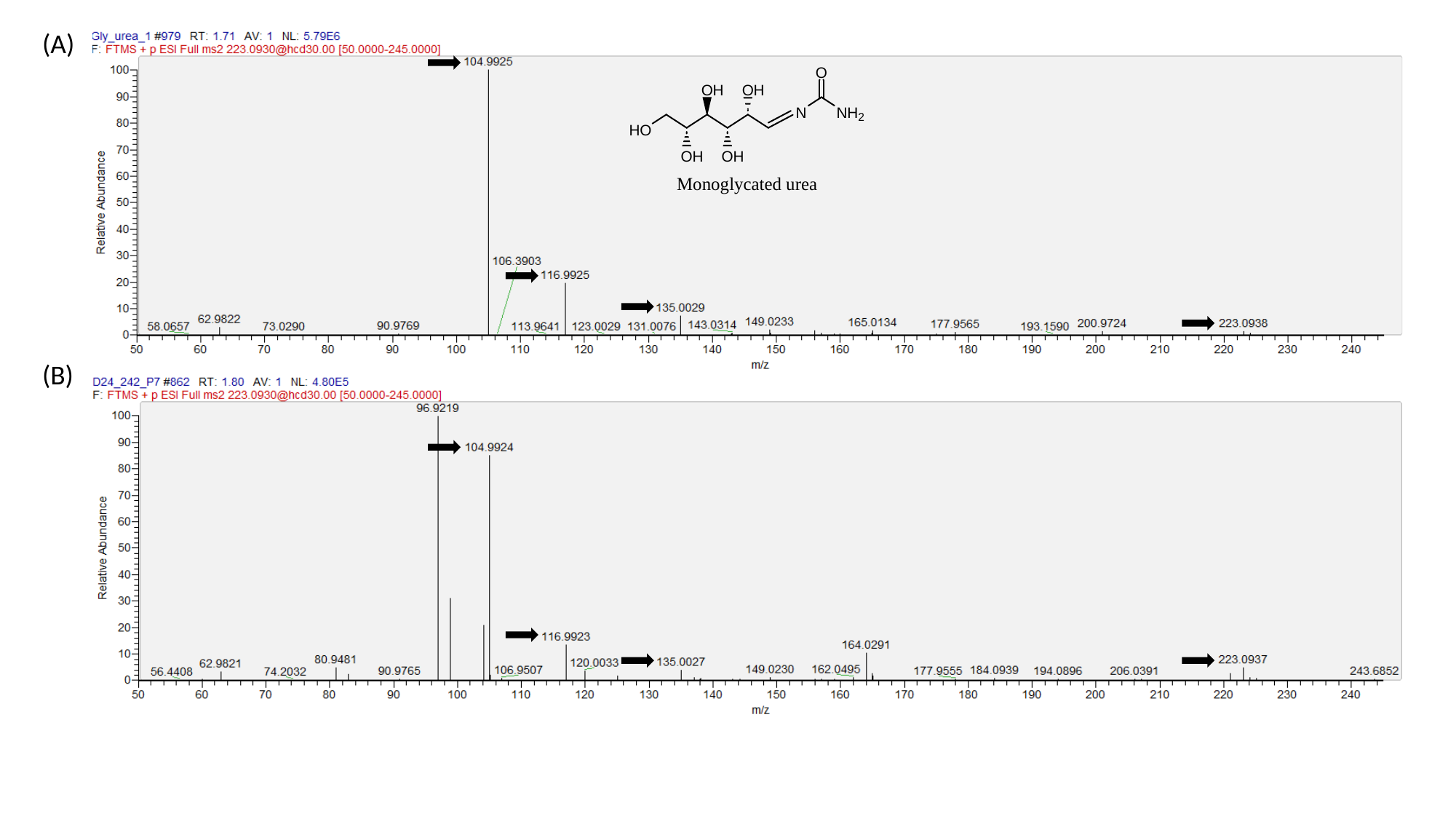

(A)
(B)

### Slide 36
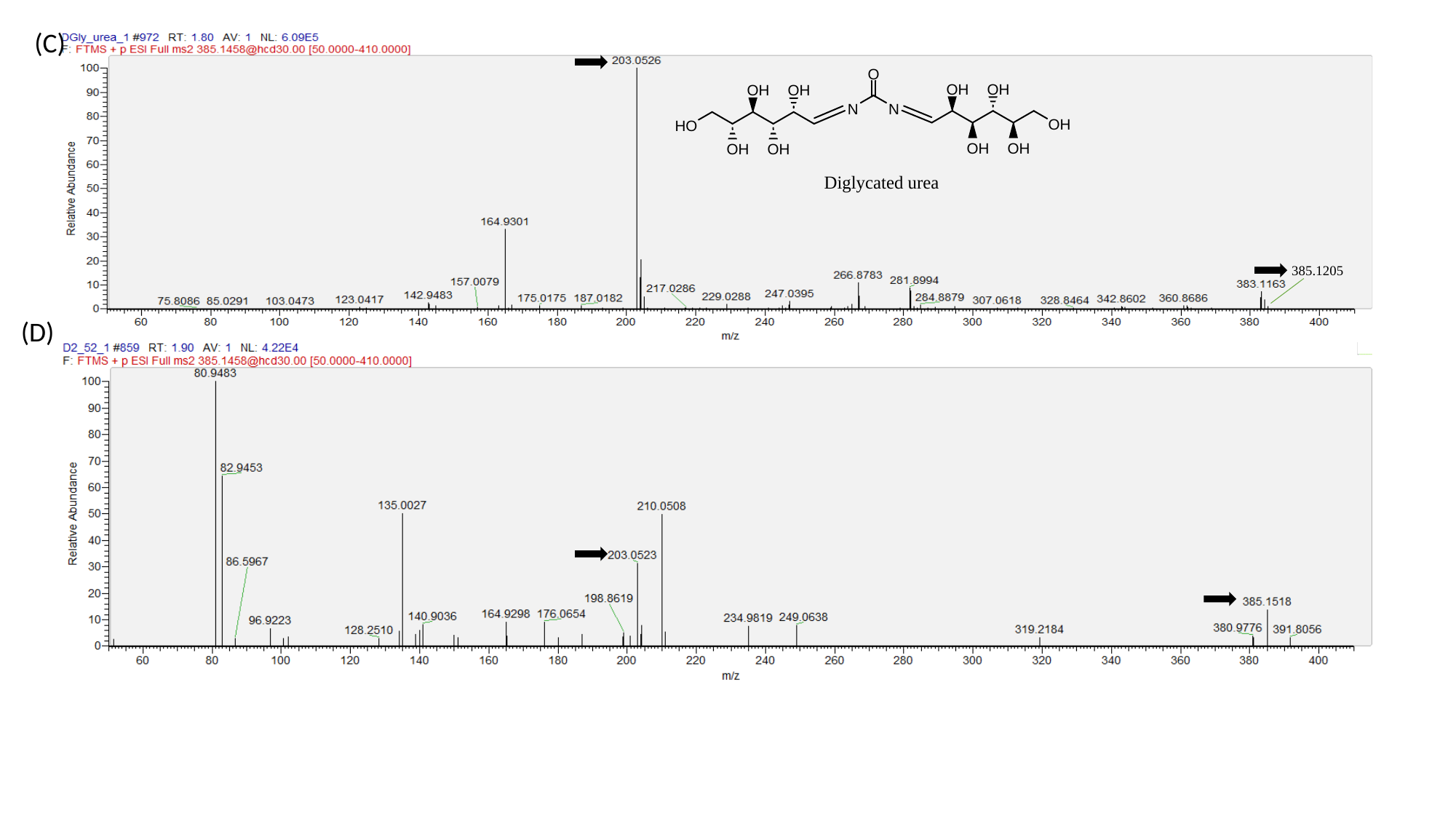

(C)
385.1205
(D)

### Slide 37
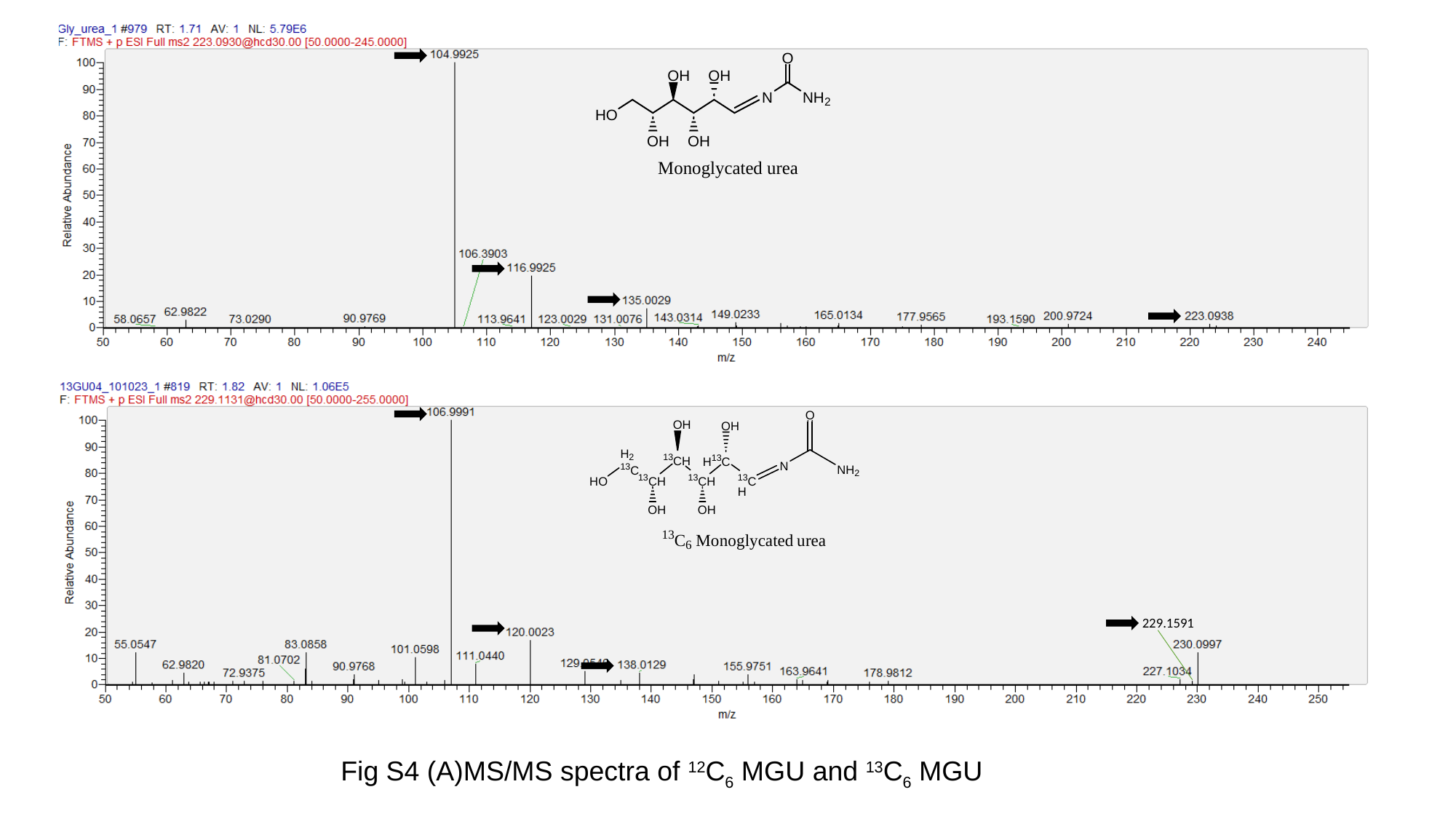

229.1591
Fig S4 (A)MS/MS spectra of ^12^C_6_ MGU and ^13^C_6_ MGU

### Slide 38
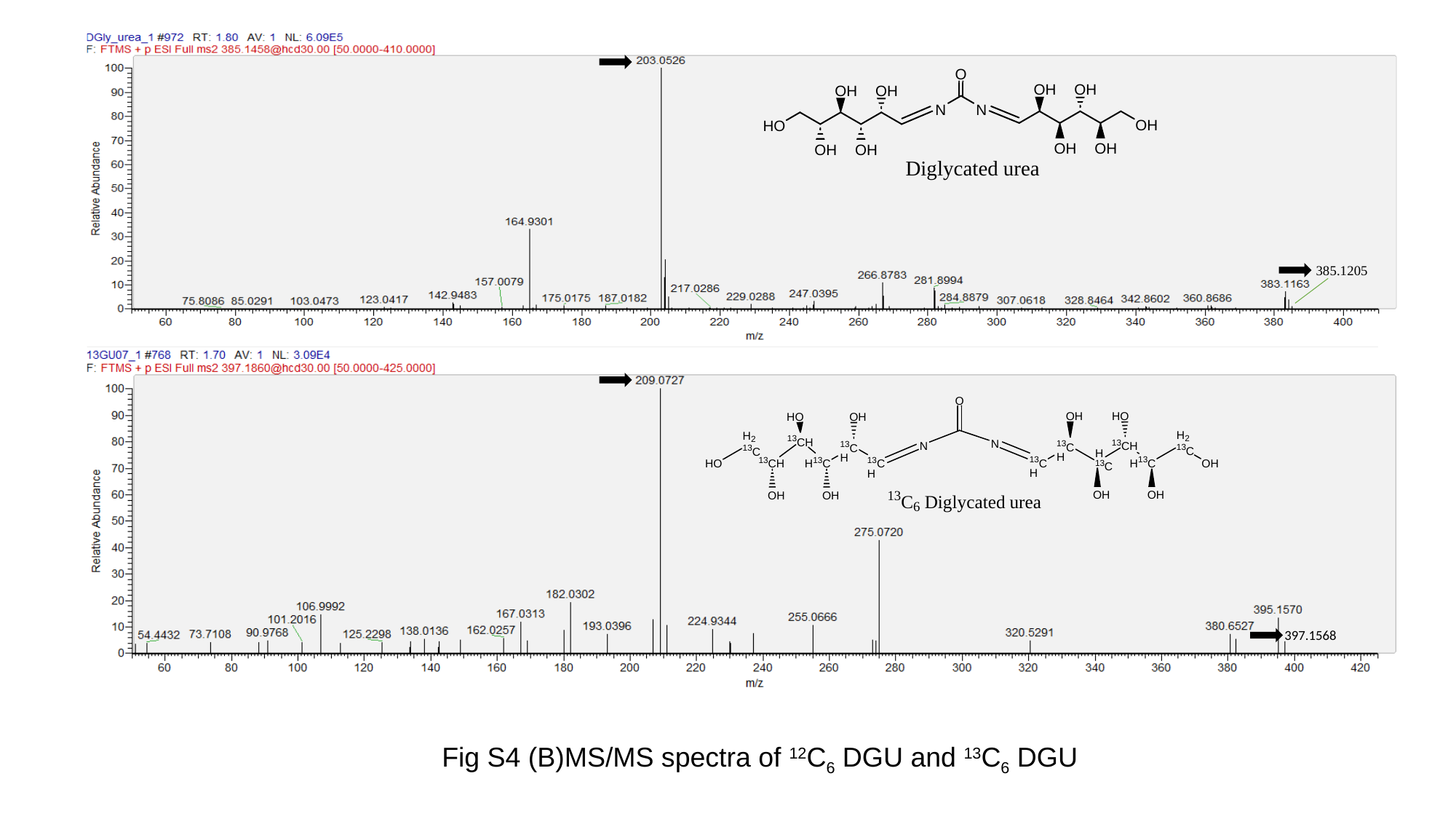

385.1205
397.1568
Fig S4 (B)MS/MS spectra of ^12^C_6_ DGU and ^13^C_6_ DGU

### Slide 39
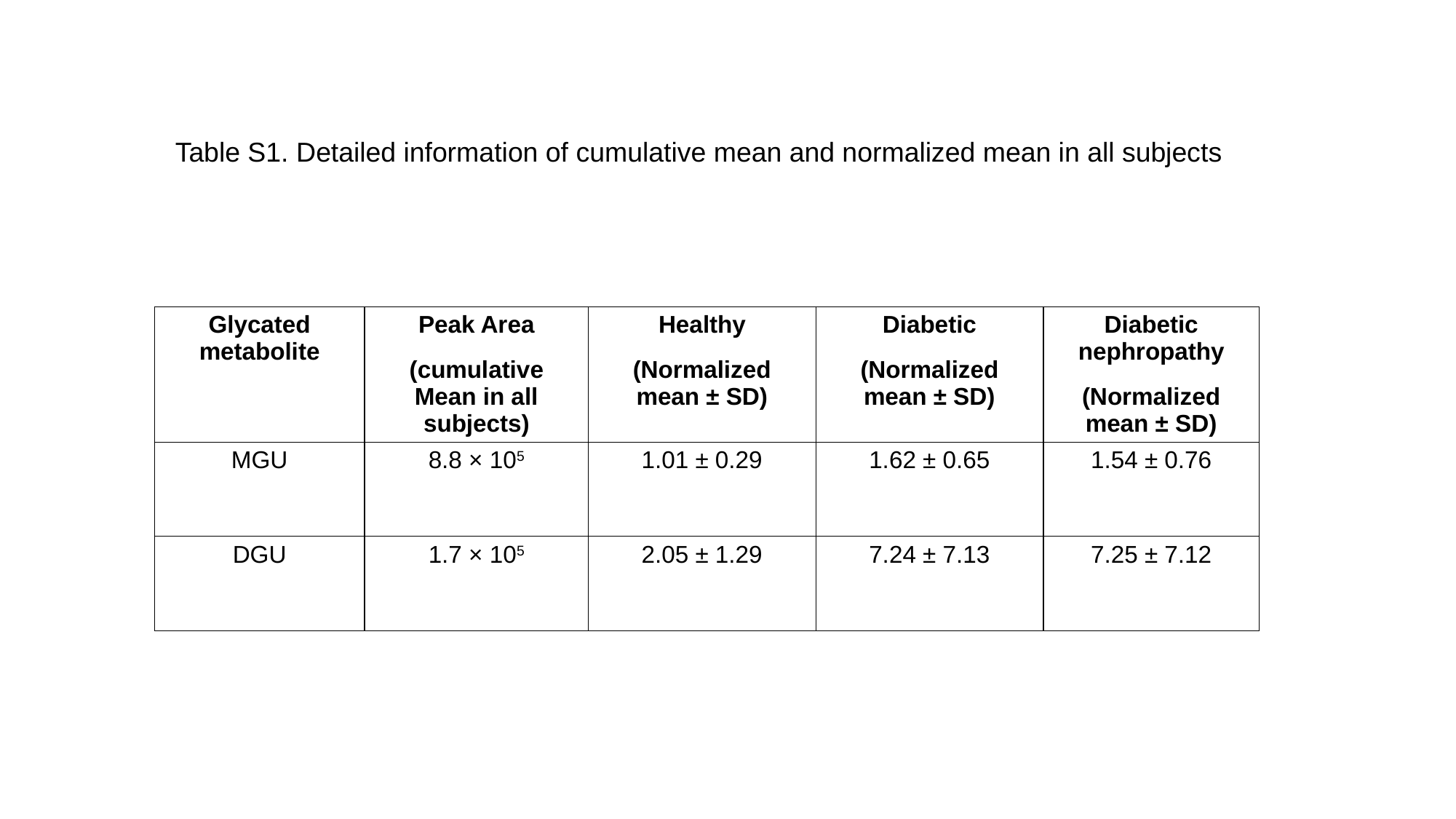

Table S1. Detailed information of cumulative mean and normalized mean in all subjects
Table S1. Detailed information of cumulative mean and normalized mean in all subjects
| Glycated metabolite | Peak Area (cumulative Mean in all subjects) | Healthy (Normalized mean ± SD) | Diabetic (Normalized mean ± SD) | Diabetic nephropathy (Normalized mean ± SD) |
| --- | --- | --- | --- | --- |
| MGU | 8.8 × 10^5^ | 1.01 ± 0.29 | 1.62 ± 0.65 | 1.54 ± 0.76 |
| DGU | 1.7 × 10^5^ | 2.05 ± 1.29 | 7.24 ± 7.13 | 7.25 ± 7.12 |
